# PARP1 recruits SPRTN to DNA-protein crosslinks through a conserved poly-ADP-ribose binding domain

**DOI:** 10.64898/2026.01.09.698661

**Authors:** Katelyn Hurley, Liam P. Leary, Martin Garcia, Ayesha Rahman, Matthew A. Schaich, Max S. Kloet, Frederick Kumi-Ansah, Elizabeth G. Blade, Qiang Liu, Andrew Rhiner, Megan Le, Quincee Simonson, Dmitri V. Filippov, Gerbrand J. van der Heden van Noort, Jason S. McLellan, Bennett Van Houten, Jaime Lopez-Mosqueda

## Abstract

DNA-protein crosslinks (DPCs) are toxic DNA lesions formed by the covalent attachment of proteins to DNA. Failure to resolve DPCs leads to genomic instability, premature aging, and cancer predisposition. Although multiple proteases and the 26S proteasome degrade DPCs, how these lesions are detected and marked for proteolysis remains unclear. Here, we show that poly-(ADP-ribose) polymerases (PARP1/2) sense DPCs and modify them with poly(ADP-ribose) (PAR) to promote repair via a SPRTN-Tdp1 axis. We discovered a Nudix homology domain (NHD) in SPRTN that mediates direct non-covalent PAR binding and is important for DPC repair. Loss of PARP1/2 activity or mutation of the SPRTN NHD leads to sustained DPCs. Single-molecule analysis revealed that SPRTN does not bind efficiently to the DPC, however after the addition of PARP1 in the presence of NAD^+^, SPRTN binding to the DPC was significantly increased. Our findings establish PARP1/2 enzymes as immediate DPC sensors, reveal PARylation as a signal marking DPCs for SPRTN-dependent degradation, and identify SPRTN as the first PARP-directed protease.

## INTRODUCTION

Genome integrity is continually challenged by endogenous and exogenous sources of DNA damage. To preserve genomic stability, cells rely on a network of surveillance and repair pathways that detect, signal, and resolve DNA lesions ^1^. These pathways are collectively called the DNA damage response (DDR), and orchestrate the precise spatial and temporal recruitment of repair factors, mediated by an array of post-translational modifications ^2, 3^. When DDR mechanisms fail, mutation and genome instability ensue, driving oncogenesis, neurodegeneration, accelerated aging, and cell death ^1^. This is exemplified by hereditary syndromes of premature aging and cancer predisposition caused by DNA repair gene mutations ^4^.

Among the diverse DNA lesions, DNA-protein crosslinks (DPCs) are particularly cytotoxic. These lesions arise when proteins become covalently attached to DNA, either through chemical exposure or aberrant enzymatic reactions. Topoisomerase I and II covalent complexes (Top1cc/Top2cc) represent well-characterized enzymatic DPCs. During their catalytic cycle, topoisomerases form transient covalent intermediates between the catalytic tyrosine and the DNA backbone. Topoisomerase poisons such as camptothecin and etoposide, exert their cytotoxicity by stabilizing these intermediates. Due to their bulk, DPCs obstruct replication, transcription, and chromatin remodeling ^5^. Persistent DPCs promote genomic instability, and cells defective in DPC repair exhibit hypersensitivity to DPC-inducing agents.

Ruijs-Aalfs syndrome, a segmental progeroid syndrome associated with genomic instability and childhood liver cancer, is caused by germline mutations in *SPRTN* ^6, 7^. SPRTN was subsequently identified as the first mammalian protease dedicated to DPC repair ^8, 9, 10^. SPRTN degrades a broad range of chromatin-associated proteins, including histones, DNA topoisomerases, and PARP1 suggesting that it must selectively target diverse substrates ^8, 9, 10^.

Effective DDR requires accurate lesion detection, followed by signaling through post-translational modifications such as phosphorylation, ubiquitylation, SUMOylation, and ADP-ribosylation, which recruit repair complexes to damage sites ^11^ ^12, 13, 14, 15^. DPCs have recently been shown to undergo ubiquitylation and SUMOylation, and disruption of these modifications hinders repair ^16, 17, 18, 19, 20, 21, 22^. However, how DPCs are selectively detected and marked for SPRTN-mediated proteolysis remains incompletely understood. Here we identify a NUDIX homology domain (NHD) in SPRTN that mediates non-covalent binding to poly(ADP-ribose) (PAR). We show that PARP1 functions as an immediate DPC sensor by PARylating Top1ccs. This modification promotes the recruitment of SPRTN and Tdp1, enabling the resolution of Top1ccs. Our findings reveal a mechanism by which SPRTN activity is regulated and broadly directed towards diverse DPC substrates, regardless of size, identity, or biochemical properties.

## RESULTS

### SPRTN and Tdp1 Recruitment to DNA damage is PARP-Dependent

SPRTN in known to localize to microirradiation-induced DNA damage sites, which harbor single- and double-strand DNA breaks, as well as DPCs ^23^. These regions are enriched with post-translational chromatin modifications including ubiquitin, SUMO, and PAR. Because PARP1 catalyzes PARylation at DNA breaks and recent work in Xenopus extracts demonstrated that PARP1 activity is important for efficient DPC resolution^24^, we hypothesized that PARP1 might function as a DPC sensor that promotes SPRTN recruitment to damage sites in mammalian cells. Mechanistically, PARylation could facilitate SPRTN accumulation either by creating a binding platform for PAR-interacting motifs within SPRTN or by remodeling chromatin to expose DPCs. To address this, we performed laser microirradiation in U2OS cells expressing GFP-SPRTN. Consistent with previous reports, GFP-SPRTN rapidly accumulated at DNA damage tracks. Strikingly, treatment with the PARP inhibitor Olaparib significantly reduced SPRTN recruitment ^24^, indicating that PARP catalytic activity is required (Figure 1A and 1B). Olaparib effectively blocked PARylation under DNA damage conditions (Figure S1A), confirming that the observed effect reflects loss of PAR synthesis. We next sought to examine SPRTN recruitment in PARP1 knockout cells^25^. Although eliminating PARP1 expression results in undetectable PAR levels (Figure S1B), SPRTN continued to localize to damage tracks in PARP1 knockout cells, most likely due to PARP2 activity (Figure S1C). To address this, we next used PARP1/2 double-knockout U2OS^26^ cells to assess whether both PARP1 and PARP2 are required for SPRTN recruitment to DNA damage. In these cells, SPRTN recruitment to DNA damage tracks was completely abolished (Figure 1C), confirming that PARP1 and PARP2 function redundantly to promote SPRTN accumulation at DNA lesions.

**Figure 1.**
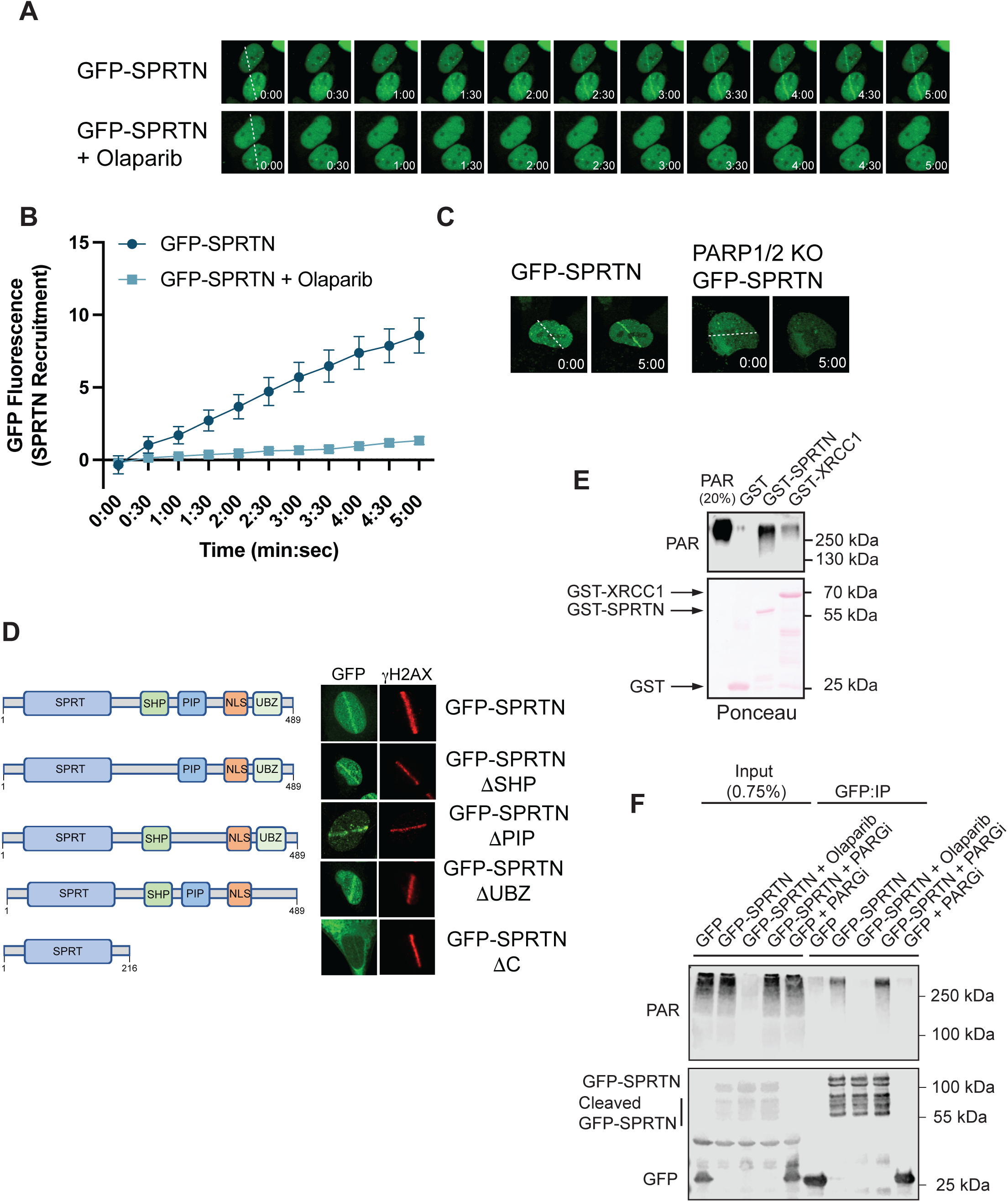
SPRTN recruitment to DNA damage is mediated by poly(ADP-ribose) (PAR). **(A)** UV-laser microirradiation of U2OS T-REx cells transiently expressing GFP-SPRTN and sensitized with BrdU. Cells were left untreated or pretreated with Olaparib (20 μM, 12 h) prior to microirradiation. Insets indicate time (min:sec); dashed lines mark the microirradiation path. **(B)** Quantification of GFP-SPRTN recruitment at damage tracks in 20 cells; error bars represent SEM. **(C)** Microirradiation of U2OS T-REx wild-type or PARP1/2 knockout cells transiently expressing GFP-SPRTN and sensitized with BrdU. Insets indicate time (min:s); dashed lines mark the microirradiation path. **(D)** Immunofluorescence of microirradiated U2OS T-REx cells stably expressing GFP-SPRTN variants. Cells were fixed immediately after microirradiation and stained for γH2AX. **(E)** GST pulldown assay using GST alone (negative control), GST-SPRTN, or GST-XRCC1 (positive control) incubated with biosynthesized PAR. Representative of three independent experiments. **(F)** GFP-SPRTN expression was induced in U2OS T-REx cells with doxycycline (1 μg/mL, 12 h). Cells were treated with Olaparib (20 μM, 12 h) to block ADP-ribosylation or PARG inhibitor (PARGi; 20 μM, 12 h) to prevent PAR degradation. GFP or GFP-SPRTN was immunoprecipitated with GFP-Trap beads and analyzed by immunoblotting. Representative of three independent experiments.

SPRTN contains three characterized protein-interacting modules in its C-terminal region – the SHP, PIP, and UBZ domains ^27, 28, 29, 30^; however, none are known to bind PAR. To test whether these domains mediate PAR-dependent recruitment, we generated GFP-SPRTN deletion mutants lacking SHP, PIP, UBZ, or the entire C-terminus (ΔC). All mutants localized to microirradiation tracks (Figure 1D), indicating that these domains are dispensable for SPRTN recruitment to DNA damage. The ΔC mutant was largely cytoplasmic, likely due to loss of the nuclear localization signal (NLS), but the fraction that entered the nucleus still recruited to damage sites. Reintroduction of the NLS into the ΔC mutant (GFP-SPRTNΔC+NLS) restored nuclear localization and robust recruitment (Figure S1D) ^10^.

We next asked whether SPRTN directly binds PAR. GST pulldown assays using biosynthesized PAR revealed that GST-SPRTN interacts with PAR in vitro, similar to XRCC1, a known PAR-binding protein (Figure 1E). To confirm this interaction in cells, we treated U2OS cells with PARG inhibitor to stabilize PAR and performed co-immunoprecipitation. SPRTN co-precipitated with PAR under these conditions (Figure 1F), demonstrating that SPRTN engages PAR in vivo. These results identify PARP activity as a critical determinant of SPRTN recruitment to DNA damage sites and reveal that SPRTN directly binds to PAR, despite lacking canonical PAR-binding motifs.

Tyrosyl-DNA phosphodiesterase 1 (TDP1), a critical enzyme for resolving Top1ccs ^31^, also localizes to DNA damage sites in a PARP-dependent manner ^32^. Moreover, a recent proteomic study identified TDP1 as a direct PAR-binding protein and mapped a region near its catalytic domain that mediates this interaction ^33^. To confirm that PAR binding is required for TDP1 recruitment, we used a wildtype TDP1 and a mutant lacking this region (TDP1_1-164_) in microirradiation assays. While wildtype TDP1 readily recruited to DNA damage tracks in a PARP-dependent manner, the TDP1 mutant exhibited impaired recruitment (Figure S1E and S1F). These results demonstrate that TDP1 recruitment depends on both PARP activity and its intrinsic PAR-binding capability. Together, these findings reveal that PARP1 regulates the recruitment of two key DPC repair enzymes, SPRTN and TDP1, through direct, non-covalent interactions with PAR.

### The SprT Domain Contains a Nudix Homology Domain That Mediates PAR Binding

Several structurally distinct motifs mediate PAR-binding ^34, 35^, but SPRTN has not been reported to harbor any canonical PAR-binding domains. Because the N-terminal half of SPRTN is sufficient for recruitment to DNA damage sites, we searched for a PAR-binding region within the SprT protease domain. Bioinformatic analysis ^36, 37^, revealed a conserved Nudix homology domain (NHD) overlapping the zinc-binding domain (ZBD) within the SprT domain (Figure 2A). NHDs are evolutionarily related to NUDIX hydrolase family, which typically contain a 23-amino acid Nudix box motif (GX_5_EX_7_REUXEEXGU, where U is a bulky aliphatic residue and X is any amino acid) ^38, 39, 40^. Unlike classical NUDIX domains, which hydrolyze substrates including PAR ^38, 39, 41, 42^, NHDs lack catalytic activity but retain PAR-binding capability.

**Figure 2.**
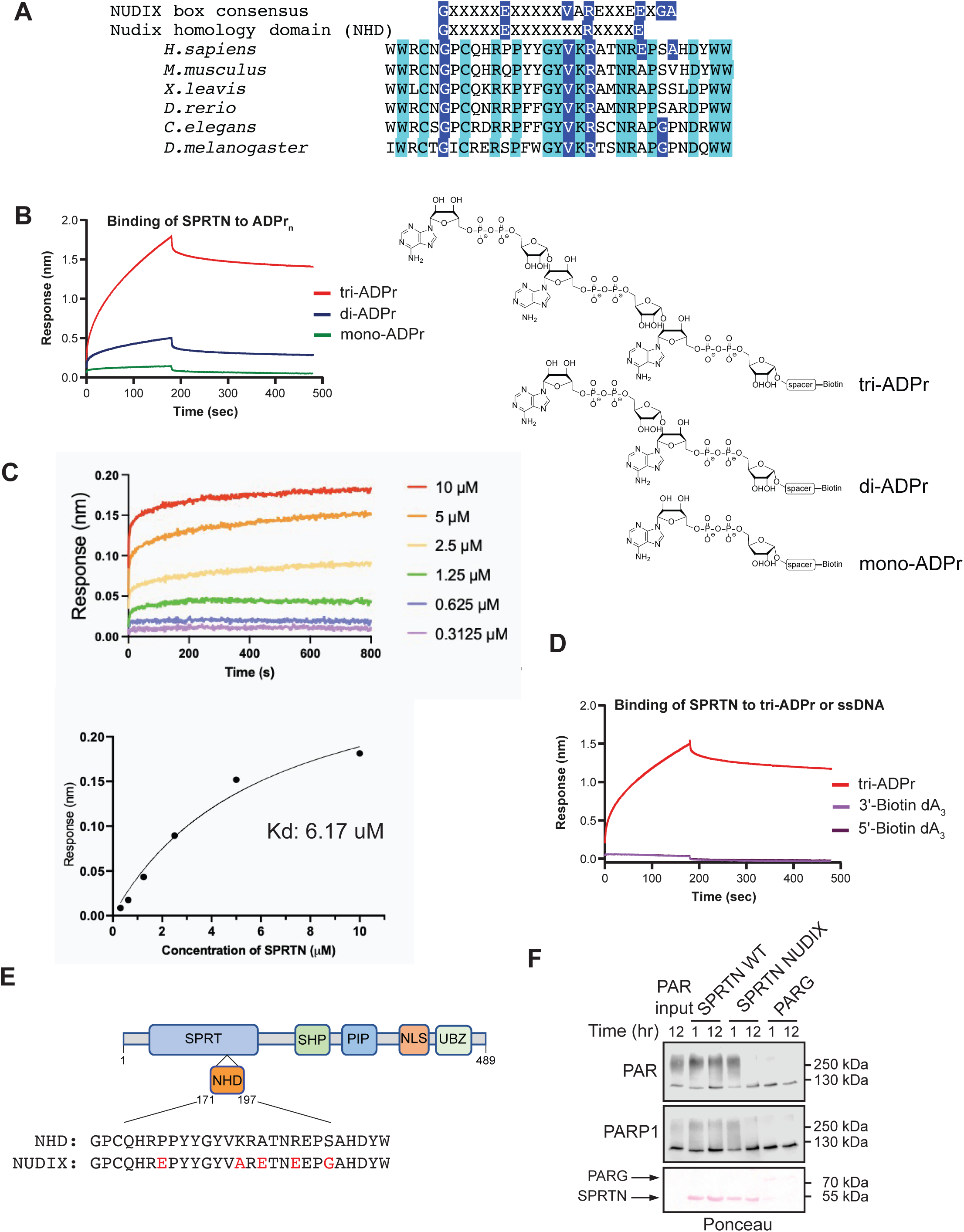
SPRTN contains a Nudix Homology Domain (NHD) that binds poly(ADP-ribose). **(A)** Sequence alignment of the SPRTN Nudix homology domain (NHD) across eukaryotic species. Conserved motifs corresponding to the canonical NUDIX box and NHD are indicated. **(B)** Biolayer interferometry (BLI) analysis of SPRTN interaction with mono-, di-, and tri-ADP-ribose (ADPr). Chemical structures of the ligands used in binding experiments are shown. **(C)** BLI sensorgrams for SPRTN binding to immobilized tri-ADPr. Increasing SPRTN concentrations were applied to measure association and dissociation kinetics. The equilibrium response was fitted to a one-site binding model to determine the dissociation constant (Kd). **(D)** Comparative BLI analysis of SPRTN binding to tri-ADPr versus biotinylated ssDNA (3′-biotin or 5′-biotin tri-deoxyadenosine). **(E)** Schematic of engineered SPRTN NHD showing five point mutations (P177E, K184A, A186E, R189E, S192G) introduced to reconstitute the canonical NUDIX motif (GX_5_EXEE). **(F)** In vitro PAR hydrolysis assay comparing wild-type SPRTN, SPRTN^NUDIX^ mutant, and PARG. Representative of three independent experiments.

The solved crystal structure of human SPRTN bound to single-strand DNA revealed a ZBD capable of interacting with nucleotides or ADP ^43^. Given the structural similarity between ADP-ribose and nucleic acids (Figure S2A), we hypothesized that the NHD mediates PAR binding. Biolayer interferometry (BLI) confirmed that SPRTN binds mono-, di-, and tri-ADP-ribose with increasing affinity, favoring extended PAR polymers (Figure 2B). To quantify this interaction, we measured the apparent dissociation constant (Kd) for tri-APD-ribose and observed a Kd of 6.17 μM using purified SPRTN (Figure 2C). Under identical conditions, we could not detect binding to a three-nucleotide ssDNA fragment (dA_3_). Orientation changes in dA_3_ did not affect binding, indicating specificity for ADP-ribose (Figure 2D) and suggesting that SPRTN preferentially recognizes ADP-ribose over nucleic acids.

To test whether SPRTN could be converted into a PAR hydrolase, we engineered a gain-of-function SPRTN^NUDIX^ mutant by restoring consensus residues within the NHD (Figure 2E). Structural modeling positioned di-ADP-ribose near glutamate 189 (E189), suggesting a potential catalytic site (Figure S2B). In vitro, SPRTN^NUDIX^ degraded PAR and PAR covalently attached to PARP1 within 12 hours, similar to PARG (Figure 2F), but did not hydrolyze RNA (Figure S2C), demonstrating specificity for PAR and indicating that wild-type SPRTN evolved for PAR binding rather than hydrolysis. Interesting, the SPRTN^NUDIX^ variant failed to bind DNA relative to wild-type SPRTN and also did not exhibit DNA hydrolysis activity (Figures S2D).

Because SPRTN undergoes self-cleavage upon binding to DNA ^8, 9, 10^, we asked whether PAR influences SPRTN cleavage. PAR alone did not trigger SPRTN cleavage. However, di- and tri-ADP-ribose inhibited DNA-dependent cleavage, suggesting that these PAR fragments compete with DNA for SPRTN binding (Figure S2E). This competitive interaction implies that PAR may act as a regulatory signal, preventing SPRTN activation in contexts where DNA binding would otherwise trigger proteolysis. Consistently, our binding assays and earlier BLI experiments (Figure 2C-D) demonstrated that SPRTN exhibits higher specificity for PAR compared to DNA. Together, these findings suggest a model in which PAR binding could sequester SPRTN away from DNA, thereby modulating its protease activity and potentially linking SPRTN function to PAR-dependent stress responses.

To assess the cellular function of the NHD, we generated SPRTN-depleted cell lines expressing doxycycline-inducible, shRNA-resistant wild-type SPRTN, the SPRTN^NHD^ mutant, and the catalytically inactive mutant SPRTN^E112A^. As expected, only wild-type SPRTN exhibited robust self-cleavage, whereas SPRTN^NHD^ and SPRTN^E112A^ mutants lacked detectable cleavage (Figure S2F). Co-immunoprecipitation showed that SPRTN^NHD^ mutant still interacts with p97/VCP, indicating that loss of cleavage is not due to misfolding (Figure S2G). Interestingly, GFP-tagged SPRTN^NHD^ displayed a cleavage pattern, likely resulting from GFP-induced dimerization or residual endogenous SPRTN activity. These findings reveal that the NHD is essential for SPRTN activation in cells, likely by mediating PAR binding. Based on these results, we propose renaming the previously annotated ZBD as the Nudix homology domain (NHD), reflecting its functional and structural similarity to PAR-binding Nudix motifs.

### Nudix Homology Domain Mediates SPRTN’s Recruitment to DNA Damage

Structural insights from the SPRTN-ADP crystal structure and di-ADP-ribose modeling suggested that the NHD could serve as a PAR-binding module critical for SPRTN function. To test this hypothesis, we introduced six-point mutations within the NHD predicted to disrupt PAR interaction (Figure 3A and 3B). GST-pulldown assays confirmed that wild-type GST-SPRTN robustly bound PAR, whereas the GST-SPRTN^NHD^ mutant did not, validating that these substitutions abolish PAR binding (Figure 3C). Importantly, the protease and NHD domains occupy opposite surfaces of SPRTN (Figure S3A), so catalytic inactivation should not influence PAR binding. Consistent with this, GST-SPRTN^E112A^ catalytic mutant retained PAR-binding capacity (Figure 3C). Mechanistically, these findings indicate that PAR binding through the NHD is not a passive interaction but a prerequisite for SPRTN’s recruitment to DNA damage sites. When we examined localization after laser microirradiation, GFP-SPRTN^NHD^ displayed markedly impaired recruitment compared to wild-type GFP-SPRTN (Figure 3D and 3E). This suggests that PAR binding acts as an early signal, positioning SPRTN at PAR-rich DNA lesions generated by PARP1/2 activity. We propose that the NHD functions as a molecular sensor for PARylation, enabling SPRTN to recognize damaged chromatin and transition into an active state for proteolysis.

**Figure 3.**
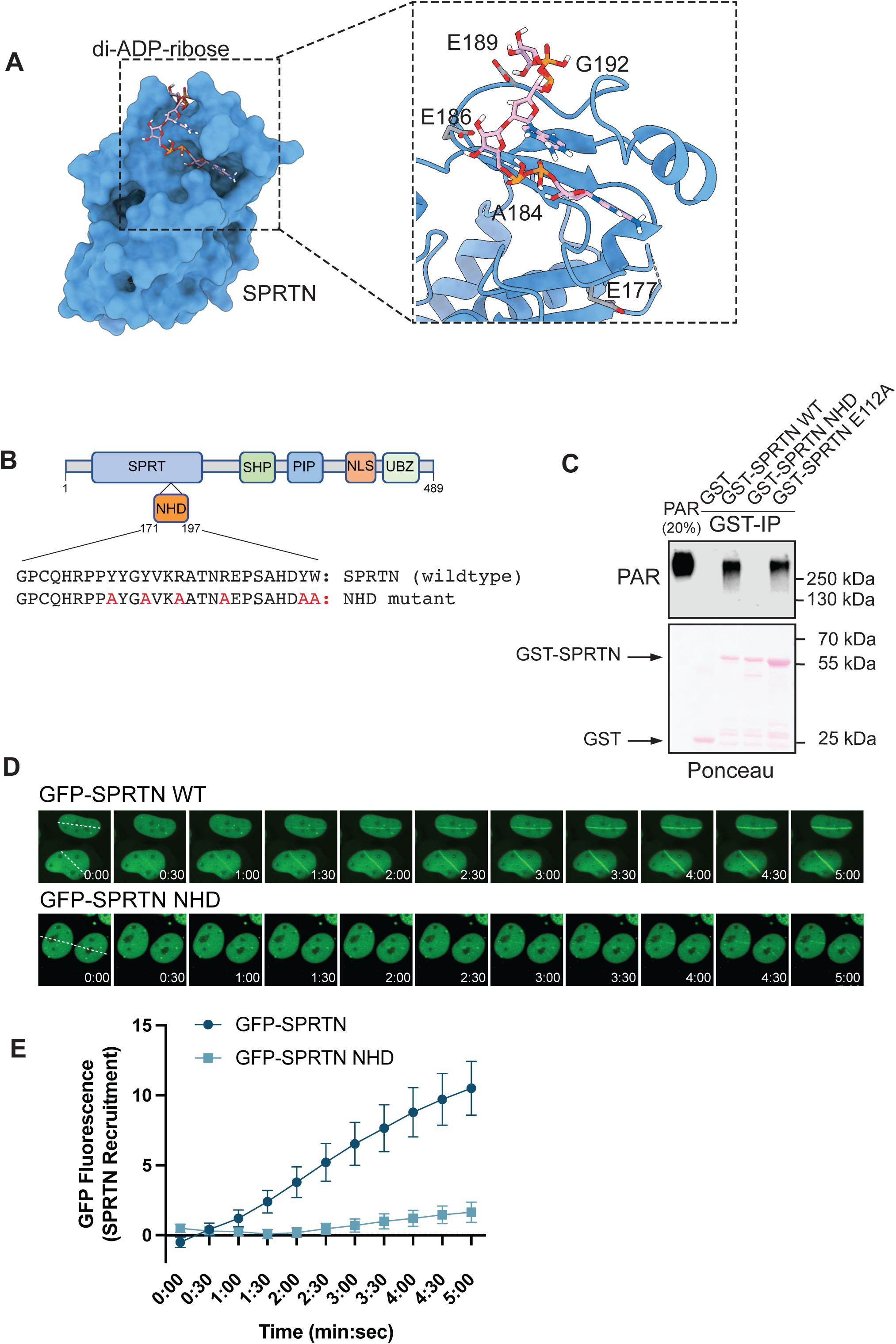
The SPRTN Nudix Homology Domain (NHD) is required for recruitment to and repair of DNA lesions. **(A)** Structural model of SPRTN (PDB: 6MDX) docked with di-ADP-ribose, illustrating the predicted PAR-binding pocket. Inset shows five candidate amino acid residues within the NHD targeted for engineering to restore the canonical NUDIX motif and enable hydrolysis of poly(ADP-ribose). **(B)** Schematic of SPRTN NHD mutations designed to abolish PAR binding. Six residues within the predicted PAR-binding interface were substituted (Y179A, Y182A, R185A, R189A, Y196A, W197A). **(C)** GST pulldown assay comparing GST alone, GST-SPRTN, GST-SPRTN^NHD^ mutant, and GST-SPRTN^E112A^ incubated with biosynthesized PAR. Representative of three independent experiments. **(D)** UV-laser microirradiation of U2OS T-REx cells transiently expressing GFP-SPRTN or GFP-SPRTN^NHD^ and sensitized with BrdU. Insets indicate time (min:s); dashed lines mark the microirradiation path. **(E)** Quantification of GFP-SPRTN and GFP-SPRTN^NHD^ recruitment at damage tracks in 20 cells; error bars represent SEM.

### Single-Molecule Analysis of PARP-Dependent SPRTN Recruitment

A central unresolved question in the field is how SPRTN recognizes the wide variety of DPCs that arise in cells. These lesions involve diverse proteins with distinct features and functions, yet SPRTN must identify and process them efficiently to maintain genome integrity. Considering the findings here, we hypothesize that SPRTN does not rely solely on intrinsic substrate features but rather responds to a damage-associated signal generated by PARP1. We propose that PARP1 can detect any DPC, regardless of its identity and use PAR to serve as a recruitment signal for SPRTN-dependent proteolysis. To directly test this model, we sought a defined DPC substrate that is not typically encountered in mammalian cells but capable of forming a covalent linkage to DNA, allowing us to study recognition in a controlled setting. Viral HUH-endonucleases, which generate covalent tyrosyl-DNA-phosphodiester bonds during rolling circle replication^44^, provide an ideal system. We selected the duck circovirus (DCV) HUH-endonuclease as a representative viral protein to create a stable, site-specific DPC in vitro and examine whether PARP1 can flag such noncanonical substrates for SPRTN recruitment. We expressed and purified DCV and confirmed its ability to form a DPC with its cognate DNA sequence (Figure 4A). Benzonase treatment reduced electrophoretic mobility, validating the covalent linkage. Similarly, treatment of the DPC with EDTA, which chelates the divalent metal ions required for DNA cleavage, confirmed the covalent nature of the linkage. Under these conditions, DCV formed a DPC with either single- or double-stranded DNA. Using this defined DPC, we next asked whether PARP1 could modify the crosslinked protein. We formed and validated DPC formation prior to adding purified PARP1 and NAD^+^ to the reactions. Remarkably, PARP1 PARylated DCV when engaged in a DPC, and this modification was inhibited by Olaparib (Figure 4B), indicating that PARP1 detects and modifies DPCs.

**Figure 4.**
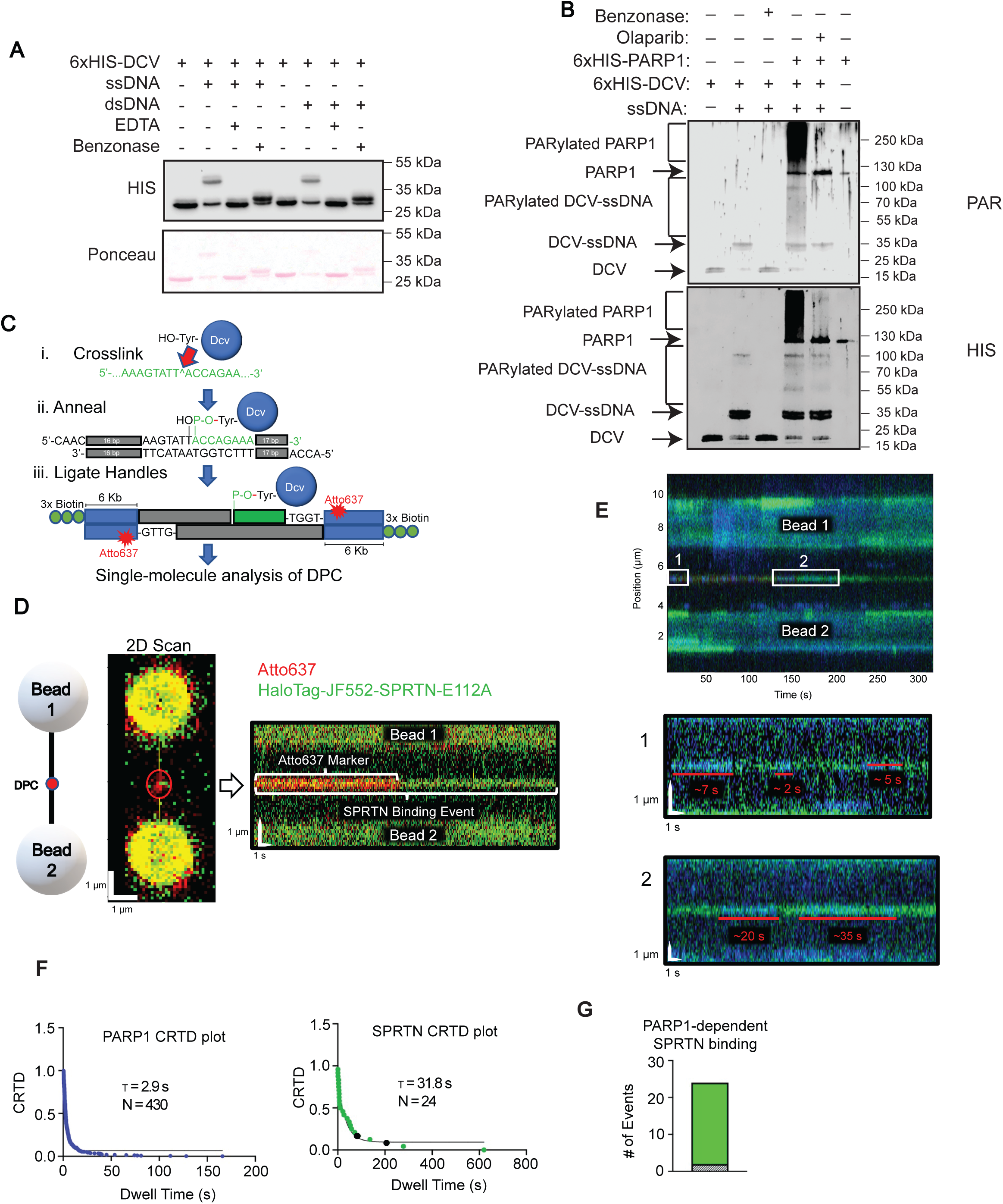
Single-molecule visualization of PARP1-dependent SPRTN recruitment to a defined DNA–protein crosslink (DPC). (A) In vitro DPC assay. Recombinant DCV (20 pmol) was incubated with single- or double-stranded DNA (10 pmol) in the absence or presence of EDTA (100 μM). Benzonase (20 units) was added to confirm DPC formation. (B) In vitro PARylation assay. DPCs were formed as in (A). Benzonase (5 units) was added to confirm DPC formation. DPC was treated with PARP1 (10 pmol) in the presence of NAD. (C) Schematic of the engineered DPC substrate for single molecule analysis. A synthetic oligonucleotide containing the duck circovirus (DCV) origin sequence (green) was covalently crosslinked to DCV HUH-endonuclease (blue sphere, ∼33 kDa) at the indicated site (red arrow). (D) Cartoon representation of the tethered DPC-containing DNA substrate labeled with Atto647 (red) and suspended between two 1.7 μm streptavidin-coated polystyrene beads. Right panels show a confocal scan of the tethered DNA (Atto647 in red) and a representative kymograph illustrating Halo-JF552–SPRTN^E112A^ binding events (magenta) relative to the DPC marker (red). Binding events appear as horizontal lines because time is plotted on the X-axis and position along the DNA on the Y-axis. (E) Representative kymographs showing sequential and overlapping binding of YFP-PARP1 (yellow) and Halo-JF552–SPRTN^E112A^ at the DPC site. Top panel (Box 1): PARP1 binds first, followed by SPRTN^E112A^; PARP1 exhibits multiple association/dissociation cycles while SPRTN remains bound. Bottom panel (Box 2): PARP1 rebinds repeatedly while SPRTN is stably engaged. Red bars indicate approximate PARP1 dwell times. (F) Cumulative residence time distribution (CRTD) plots for PARP1 (blue, N = 430 events) and SPRTN^E112A^ (green, N = 24 events) fitted to single-exponential decay functions. Mean lifetimes: PARP1 = 2.9 s; SPRTN^E112A^ = 31.8 s. Black points on the SPRTN plot denote events not preceded by PARP1 binding. (G) Quantification of SPRTN binding events relative to PARP1 occupancy. Of 24 SPRTN events, 22 (92%) occurred after or during PARP1 binding (green), whereas only 2 (8%) occurred independently (striped). These data indicate that SPRTN recruitment to DPCs is strongly PARP1-dependent.

Building on this working model, we sought to visualize DPC recognition at single-molecule resolution using the LUMICKS C-Trap optical tweezers system. To achieve this, we generated an oligonucleotide containing the DCV ori sequence, annealed to the other strand and ligated the duplex into 6 kb biotinylated handles containing Atto637 fluorophores flanking the DPC to mark its location (Figure 4C and Figure S3B). YFP-PARP1, previously shown to retain catalytic activity, and HaloTag-SPRTN^E112A^ (a catalytically inactive mutant to prevent self-cleavage) was fluorescently labeled with Janelia Fluor 552 HaloTag ligand. After confirming DPC formation in vitro (Figure S3C), we monitored protein binding in real time using kymographs (Figure 4D). Flowing HaloTag-SPRTN^E112A^ alone through the C-Trap system resulted in minimal association with the DPC. In contrast, YFP-PARP1 rapidly colocalized to the DPC site, producing numerous transient binding events. When PARP1 and NAD^+^ were introduced prior to SPRTN, we observed robust and prolonged SPRTN recruitment to the DPC (Figure 4D and 4E). Quantitative analysis of cumulative residence time distributions (CRTD) revealed that PARP1 exhibited a mean lifetime of 2.9 s (N = 430 events), whereas SPRTN^E112A^ displayed a markedly longer mean lifetime of 31.8 s (N = 24 events) (Figure 4F). Importantly, 92% of SPRTN binding events occurred after or during PARP1 occupancy, while only 8% were independent of PARP1 (Figure 4G). These single-molecule observations provide direct mechanistic evidence that PARP activity facilitates SPRTN recruitment, reinforcing our model that PARylation serves as a critical signal for DPC proteolysis. By engineering a defined DPC with a viral HUH-endonuclease and visualizing recruitment dynamics under controlled DNA tension, we demonstrate that SPRTN engagement at DPC sites depends on PARP1 and extends beyond canonical topoisomerase-derived lesions to diverse protein substrates.

### PARP1 Regulates SPRTN-Dependent DPC Proteolysis

We next aimed to understand the functional relevance of SPRTN-PAR binding with respect to DPC repair. To monitor Topoisomerase 1 covalent complexes (Top1cc), we used an antibody that specifically recognizes the tyrosyl-DNA phosphodiester linkage formed when topoisomerase 1 is covalently attached to DNA ^45^. Camptothecin (CPT) stabilizes Top1cc by blocking the re-ligation step ^46^, enabling robust formation of these lesions. Using this assay, we first tested whether SPRTN’s NHD domain and catalytic activity are required for Top1cc resolution. Endogenous SPRTN was depleted by shRNA (Figure S4A), and cells were reconstituted with shRNA-resistant wildtype SPRTN, SPRTN^NHD^ mutant, or SPRTN^E112A^ catalytic mutant (Figure S4B). Following CPT treatment and recovery, more than 60% of cells expressing wildtype SPRTN cleared Top1cc, whereas only 9% of cells expressing the SPRTN^NHD^ mutant recovered, which was similar to cells expressing SPRTN^E112A^ mutant (Figure 5A and 5B). These results indicated that PAR binding via the NHD is essential for Top1cc resolution. To determine whether PAR binding or DNA binding is more critical for DPC repair, we attempted to generate separation-of-function mutants. Our goal was to uncouple these activities and assess their individual contributions to DPC resolution. We engineered SPRTN^DNA^, which we predicted to lack DNA binding but retain PAR binding, and SPRTN^PAR^, which cannot bind PAR but binds DNA (Figure S4C). Both mutants were expressed in SPRTN-depleted cells and tested for Top1cc resolution. Strikingly, both mutants were compromised in resolving Top1cc compared to cells expressing wildtype SPRTN (Figure 5C), demonstrating that SPRTN binding to both PAR and DNA binding are essential for efficient DPC repair. Finally, we examined the role of PARP1/2 depleted cells in Top1cc resolution. CPT treatment induced Top1cc in both control and PARP1/2 double-knockout cells; however, after a 1-hour recovery, 12% of control cells retained Top1cc compared to 80% of PARP-deficient cells (Figure 5D and 5E). These findings confirm that PARP1/2-mediated PARylation is critical for Top1cc removal and acts in concert with SPRTN’s PAR- and DNA-binding activities. Together, these results establish that SPRTN-mediated proteolysis of Top1cc requires dual engagement of PAR and DNA and depends on PARP1/2 activity, revealing a coordinated mechanism for DPC resolution.

**Figure 5.**
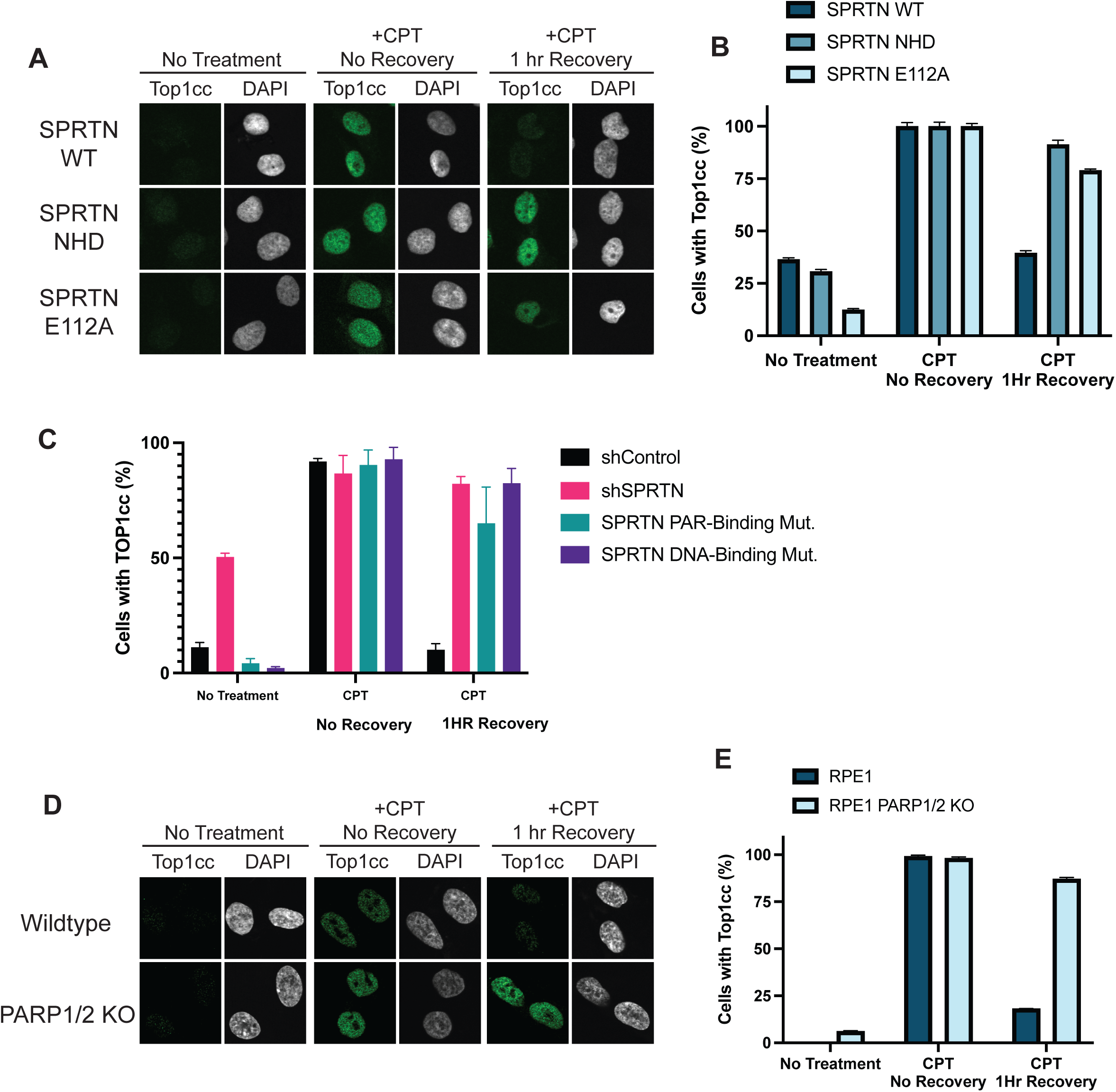
PARP1 deficiency leads to accumulation of Topoisomerase 1 covalent complexes (Top1cc). (A) Immunofluorescence detection of Top1cc in U2OS T-REx cells expressing shRNA-resistant SPRTN variants (wild-type, NHD mutant, or E112A) in an shRNA SPRTN background. Cells were untreated, treated with CPT (10 μM, 1 h), or treated with CPT followed by recovery in drug-free media for 1 h. (B) Quantification of Top1cc from Figure 5A. Graph represents scoring of 100 cells per genotype and treatment, across three independent experiments; error bars indicate SEM. (C) Quantification of Top1cc in cells expressing shRNA control, shRNA SPRTN, shRNA resistant SPRTN^PAR^ or SPRTN^DNA^. Cells were untreated, treated with camptothecin (CPT; 10 μM, 1 h), or treated with CPT followed by recovery in drug-free media for 1 h. Graph represents 100 cells per genotype and treatment across three independent experiments; error bars indicate SEM. (D) Immunofluorescence detection of Top1cc in RPE1-hTERT wild-type and PARP1/2 double-knockout cells. Cells were untreated, treated with camptothecin (CPT; 10 μM, 1 h), or treated with CPT followed by recovery in drug-free media for 1 h. (E) Quantification of Top1cc from Figure 5D. Graph represents scoring of 100 cells per genotype across three independent experiments; error bars indicate SEM.

### SPRTN and PARP1 Co-expression in Yeast Rescues CPT sensitivity in wss1Δtdp1Δ Cells

SPRTN is conserved among metazoans, whereas simple eukaryotes such as *Saccharomyces cerevisiae* express a related zinc metalloprotease, Wss1 ^47, 48^. Like SPRTN, Wss1 contains a HEXXH motif conferring proteolytic activity and able to recognize and degrade DPCs, but it harbors distinct interaction modules, including SUMO-interacting motifs (SIMs). In yeast, Wss1 and Tdp1 act together to resolve Top1cc ^49^. Accordingly, *wss1Δtdp1Δ* double mutants exhibit marked sensitivity to CPT ^47, 48^.

In a previous study, we tested whether heterologous SPRTN expression could rescue CPT sensitivity in *wss1Δtdp1Δ* cells and observed no rescue, suggesting that Wss1 and SPRTN recognize DPCs through distinct mechanisms ^10^. Given our findings that PARP-dependent PARylation earmarks DPCs for SPRTN degradation, we revisited this question by co-expressing SPRTN and PARP1 in *wss1Δtdp1Δ* cells. Yeast lack endogenous PARP1, and its heterologous expression reduces cellular fitness^50^. To circumvent suppressor mutations that arise in chronic Wss1 and Tdp1-deficient strains, we employed a doxycycline-inducible Wss1 degron allele to acutely deplete Wss1 in *tdp1Δ* cells upon doxycycline treatment ^51^ (Figure S5A). As expected, overexpression of SPRTN or PARP1 alone did not confer CPT tolerance, indicating that PARP1 expression alone does not enhance survival under DPC-inducing agents (Figure 6A and Figure S5B). Conversely, combined expression of SPRTN and PARP1 partially rescued CPT-induced lethality in *wss1Δtdp1Δ* cells (Figure 6A). Importantly, co-expression of PARP1 with SPRTN^NHD^ mutant or SPRTN^E112A^ failed to rescue CPT sensitivity, demonstrating that SPRTN’s proteolytic activity and PAR-binding capability are both essential for complementing Wss1 function. These results provide in vivo evidence that SPRTN-mediated proteolysis of DPCs is regulated by PARP1 and that PARylation serves as a critical signal for SPRTN function. Our yeast reconstitution system highlights the evolutionary divergence of DPC repair pathways and underscores the requirement for PARP-SPRTN cooperation in metazoan cells.

**Figure 6.**
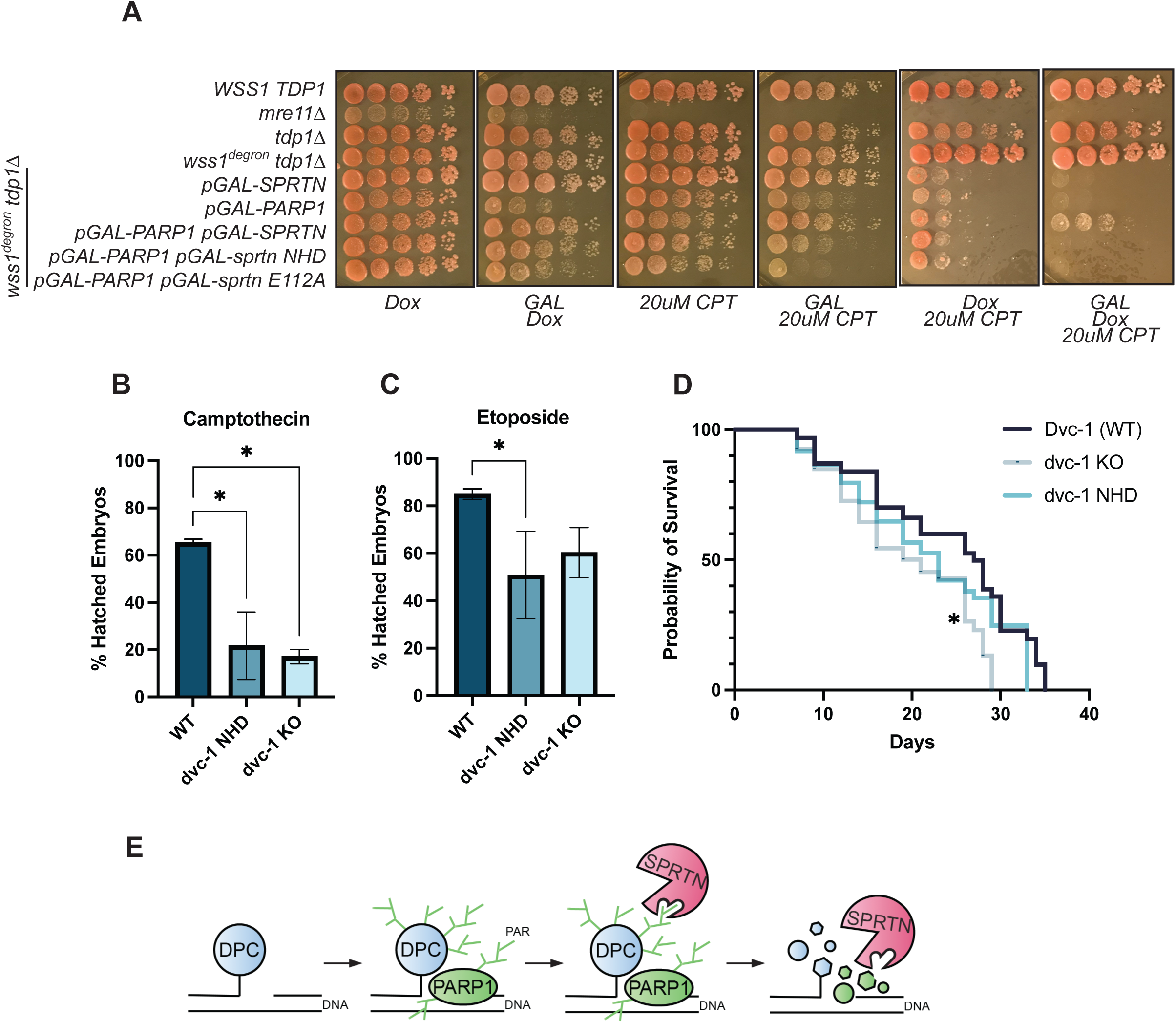
PARP1 and SPRTN co-expression rescues DPC repair defects in yeast, and dvc-1 NHD mutation sensitizes *C. elegans* to DPC-inducing drugs. **(A)** Yeast spot assay showing five-fold serial dilutions of indicated genotypes. Cells were plated on YEP medium containing 2% galactose or glucose, 10 μM doxycycline, and/or 20 μM camptothecin (CPT). Plates were incubated for 3 days at 30 °C. Representative of three independent experiments. **(B, C)** Hatching rates of *C. elegans* following exposure to CPT (1 μM; B) or etoposide (50 μM; C). Representative of three independent experiments; error bars indicate SEM; *p < 0.05. **(D)** Longevity analysis of *C. elegans* expressing wild-type Dvc-1, dvc-1^NHD^ mutant, or dvc-1 knockout. Lifespan was measured in n = 60 worms per genotype across two independent experiments. A statistically significant reduction in lifespan was observed in dvc-1 knockout animals compared with wild type (p = 0.0075, Mantel–Cox test). **(E)** Working model illustrating PARP1-dependent regulation of SPRTN-mediated DPC repair.

### SPRTN–PAR Interaction is Essential for DPC Tolerance in Multicellular Organisms

To validate the physiological importance of SPRTN-PAR interaction, we engineered a NHD mutant of dvc-1 (the SPRTN ortholog in *C.elegans*) using CRISPR-Cas9. Three genotypes were analyzed: N2 (wildtype), dvc-1^NHD^ mutant, and dvc-1 knockout (Figure S5C). Previous work showed that dvc-1 knockout embryos are hypersensitive to CPT and etoposide, which stabilize Top1cc and Top2cc, respectively ^52^. We therefore asked whether *dvc-1^NHD^* mutant embryos exhibit similar sensitivity. Following CPT exposure, over 60% of wildtype embryos survived, whereas only 20% of *dvc-1^NHD^* and 17% of *dvc-1* knockout embryos survived (Figure 6B). Likewise, etoposide treatment resulted in 85% survival for wildtype but only 50% of *dvc-1^NHD^* and 60% of *dvc-1* knockout embryos (Figure 6C). These findings demonstrate that PAR binding by SPRTN is critical for tolerance to Top1cc and Top2cc. Given that hypomorphic SPRTN mutations cause segmental progeria in humans, we next examined whether the loss of PAR binding impacts organismal longevity. Lifespan analysis revealed that only *dvc-1* knockout worms displayed reduced longevity, whereas *dvc-1^NHD^* worms aged normally (Figure 6D), suggesting that while the NHD—and thus the SPRTN–PAR interaction—is essential for embryonic viability, it is not required for lifespan maintenance in adults. Together, these results establish that SPRTN’s ability to bind PAR is essential for multicellular organisms to withstand topoisomerase-induced DPCs and maintain longevity, highlighting the physiological relevance of PARP-regulated proteolysis beyond cellular models.

## DISCUSSION

DNA–protein crosslinks (DPCs) represent one of the most structurally diverse and cytotoxic DNA lesions, yet how cells detect and mark these adducts for proteolytic removal has remained unclear. Here, we uncover a previously unrecognized role for PARP1/2 in orchestrating DPC repair through a SPRTN–Tdp1 axis (Figure 6E). Our data demonstrate that PARP1/2-dependent PARylation acts as a biochemical signal that earmarks DPCs for SPRTN-mediated degradation. We identify a Nudix homology domain (NHD) within SPRTN that enables non-covalent interaction with PAR, a property essential for SPRTN recruitment and proteolytic activation. This region, previously annotated as a zinc-binding domain (ZBD) for single-stranded DNA binding, preferentially binds tri-ADP-ribose over ssDNA. Moreover, engineering the NHD into a canonical Nudix domain converts SPRTN into a PAR hydrolase without detectable activity toward DNA or RNA, underscoring its evolutionary specialization for PAR recognition.

Our findings complement and extend recent work in *Xenopus* egg extracts showing that PARP1 activity is required for efficient DPC resolution^24^. Together, these studies converge on a model in which PARylation serves as a primary signal for protease recruitment at DPC sites. While Tdp1 also binds PAR directly, the precise domain mediating this interaction remains to be defined. Nevertheless, our cellular and organismal data establish that SPRTN’s NHD is indispensable for timely clearance of Top1cc, positioning PARP1/2 as central regulators of proteolytic DPC repair. To further test the mechanism uncovered in this study, we revisited budding yeast as a heterologous system. We previously reported that SPRTN cannot functionally complement Wss1 in *Saccharomyces cerevisiae*^10^ - deletion of both *WSS1* and *TDP1*, followed by expression of SPRTN, resulted in strong hypersensitivity to CPT, consistent with the essential role of Wss1 and Tdp1 in repairing Top1cc-derived DPCs. However, in light of our finding that PARP1 recruits SPRTN to DPCs through PARylation and that SPRTN contains an NHD capable of directly binding PAR, we reasoned that SPRTN may have failed to complement Wss1 in yeast because yeast lack PARP enzymes altogether. This provided an ideal context to test whether introduction of PARP1 is sufficient to enable SPRTN-dependent DPC repair. Indeed, while SPRTN expression alone did not rescue CPT sensitivity in *wss1Δ tdp1Δ* cells, co-expression of human PARP1 and SPRTN partially restored viability, whereas SPRTN mutants defective in PAR binding or protease activity failed to complement. These findings support a model in which PARP1-mediated PARylation of DPCs is necessary to recruit SPRTN for proteolytic degradation, a mechanism further reinforced by our single-molecule analysis showing that SPRTN engages DPCs only after PARP1 modifies the lesion with PAR.

DPC repair is further complicated by the interplay of multiple post-translational modifications. In addition to PARylation, DPCs are modified by ubiquitin and SUMO chains, which recruit E3 ligases such as TRAIP and RNF4 for proteasomal degradation (Larsen et al., 2018; Liu et al., 2021). SUMO E3 ligase ZATT (ZNF451) modifies Top2cc to facilitate repair (Schellenberg et al., 2017). How these ligases recognize DPCs remains an open question. BRCA1 and Cullin 3 complexes have been shown to directly bind Top1cc (Xu et al., 2002; Bunch et al., 2021), but additional sensors likely exist. Intriguingly, RNF146 (Iduna) is a PAR-dependent ubiquitin ligase that binds PAR via its WWE domain and functions in DNA repair (Andrabi et al., 2011; Levaot et al., 2011; Kang et al., 2011). It is tempting to speculate that PARylation may act as a master signal, not only recruiting SPRTN but also engaging PAR-dependent ubiquitin ligases to coordinate proteasomal and protease-mediated DPC clearance.

In summary, our work reveals SPRTN as the first PARP-directed protease and establishes PARP1/2 as versatile regulators of genome stability. By coupling PARylation to proteolytic activity, cells deploy an adaptable system for maintaining proteostasis at DNA lesions. These findings expand the functional repertoire of PARP enzymes beyond canonical roles in single- and double-strand break repair and highlight PARylation as a multifaceted signal for resolving complex DNA–protein adducts.

## ACKNOWLEDGEMENTS

This work was supported by the grants from the South Dakota Governor’s Office of Economic Development (SD-GOED), the South Dakota Agricultural Experiment Station and The National Science Foundation (MCB-23407189). BVH acknowledges the support of NIEHS, NIH (Grant number, R35ES039638) GJvdHvN and MSK acknowledge funding from the Dutch Research Council (NWO) (grant VI.VIDI.192.011). We thank David Toczyski and Koraljka Husnjak for comments on the manuscript, Shan Zha for U2OS PARP1 knockout cells, Keith Caldecott for the RPE1 PARP1 PARP2 double knockout cells, Nicholas Lankin for the U2OS PARP1, PARP2 and PARP1/PARP2 double knock-out cells. Didier Trono and Susan Lindquist for sharing plasmids, and Joy Scaria for the use of his *C. elegans* facility.

## AUTHOR CONTRIBUTION

J.LM. conceived the project; K.H. and J.LM. conducted the experiments with assistance from M.G., A.R., F.KA., A.R, M.L., and Q.S. L.P.L, M.S., and B.V.H performed and analyzed single molecule experiments on the C-TRAP. D.V.F and Q.L. synthesized di- and tri-ADP-ribose polymers and G.J.V. and M.S.K completed the BLI analysis. E.B and J.S.M measured affinity for tri-ADP-ribose. K.H and J.LM wrote the manuscript.

## Materials and Methods

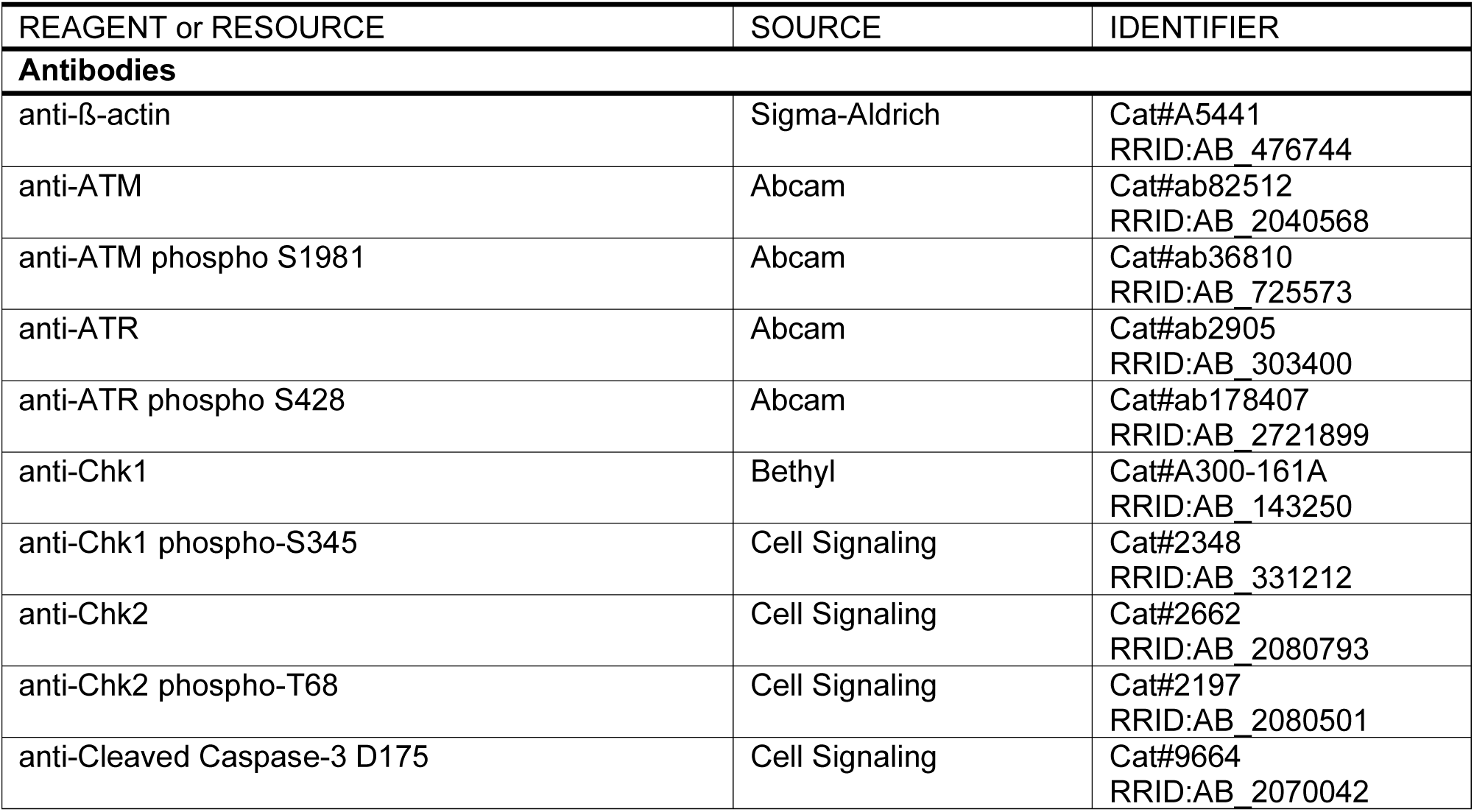
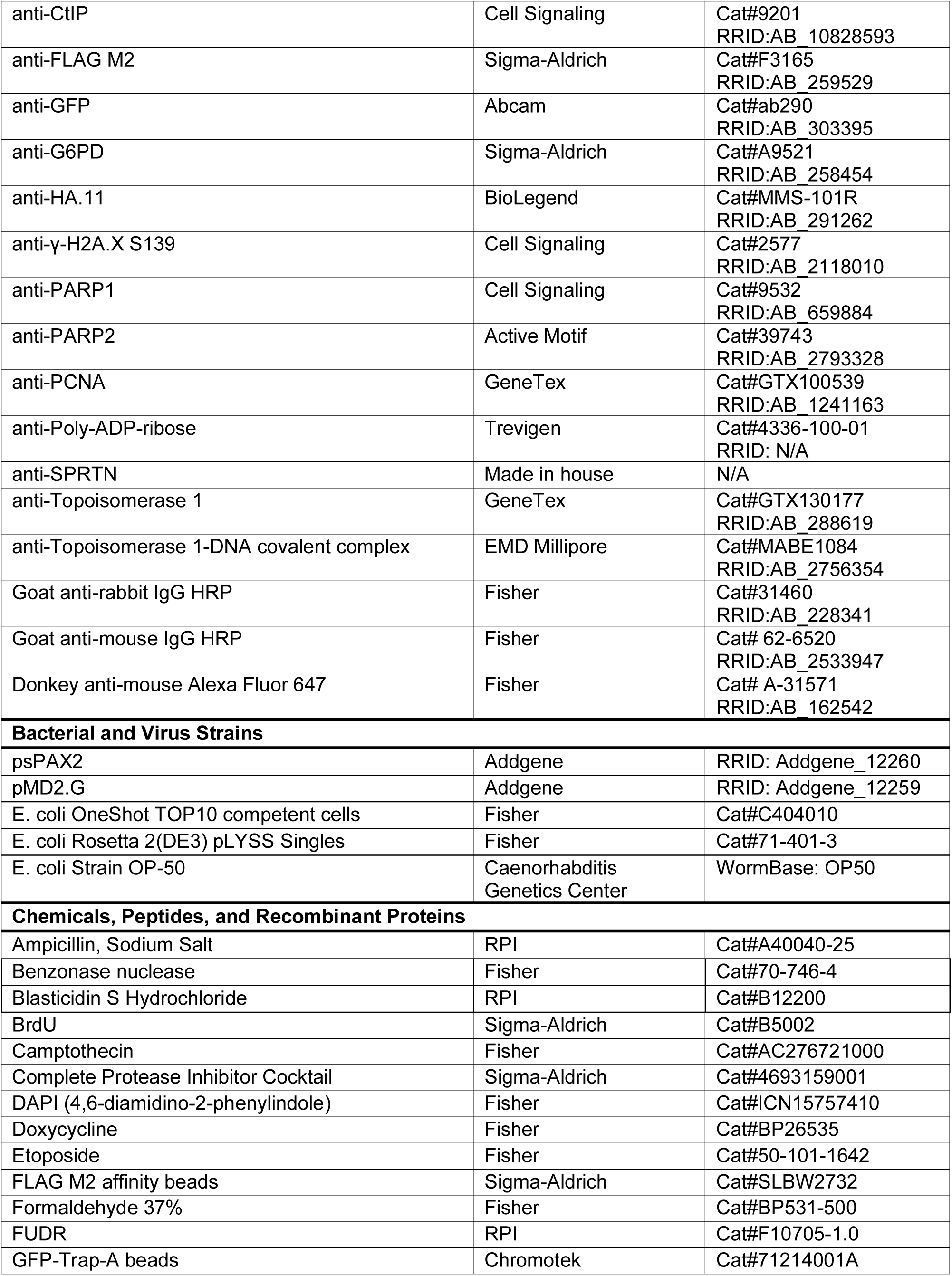
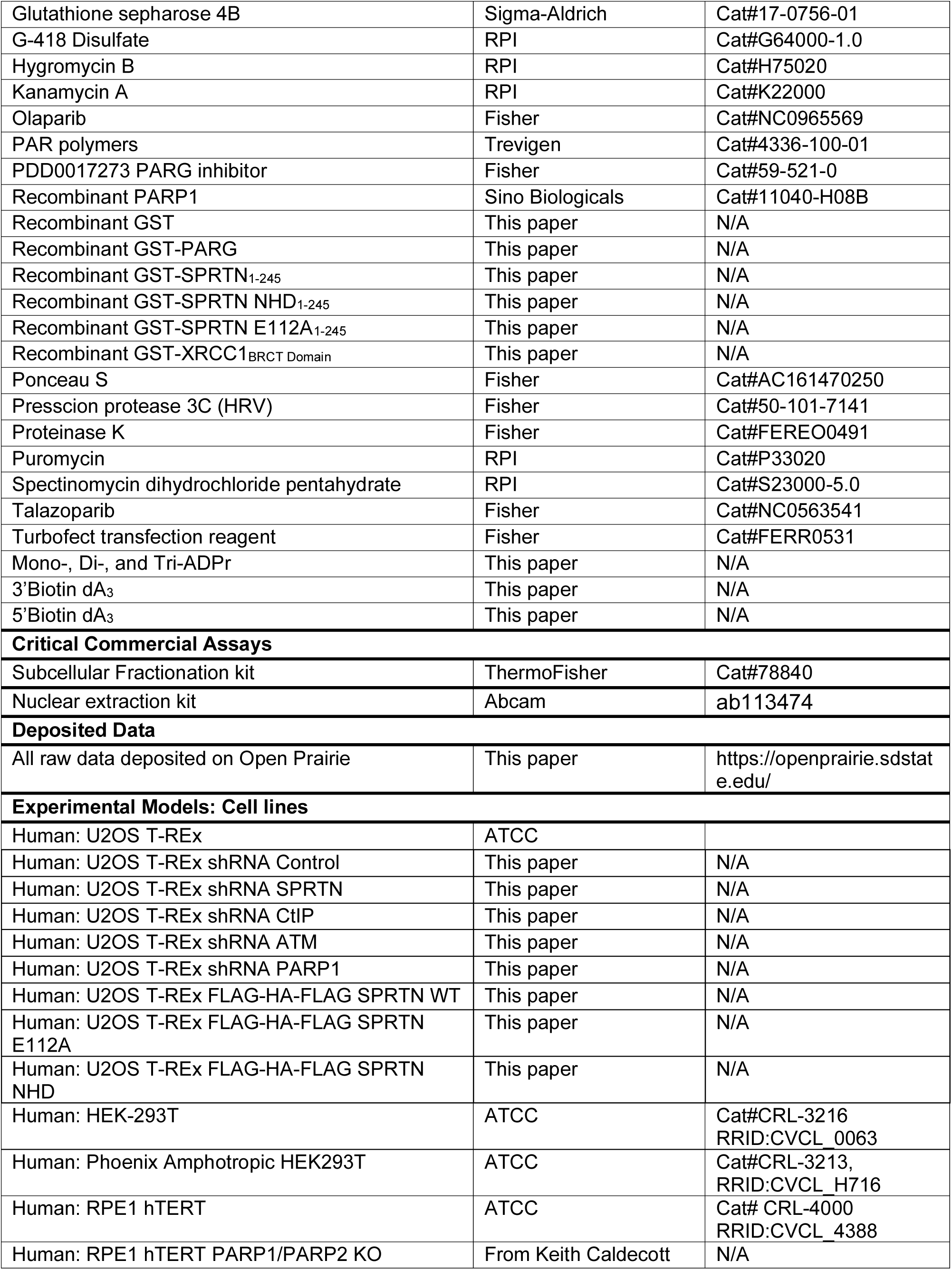
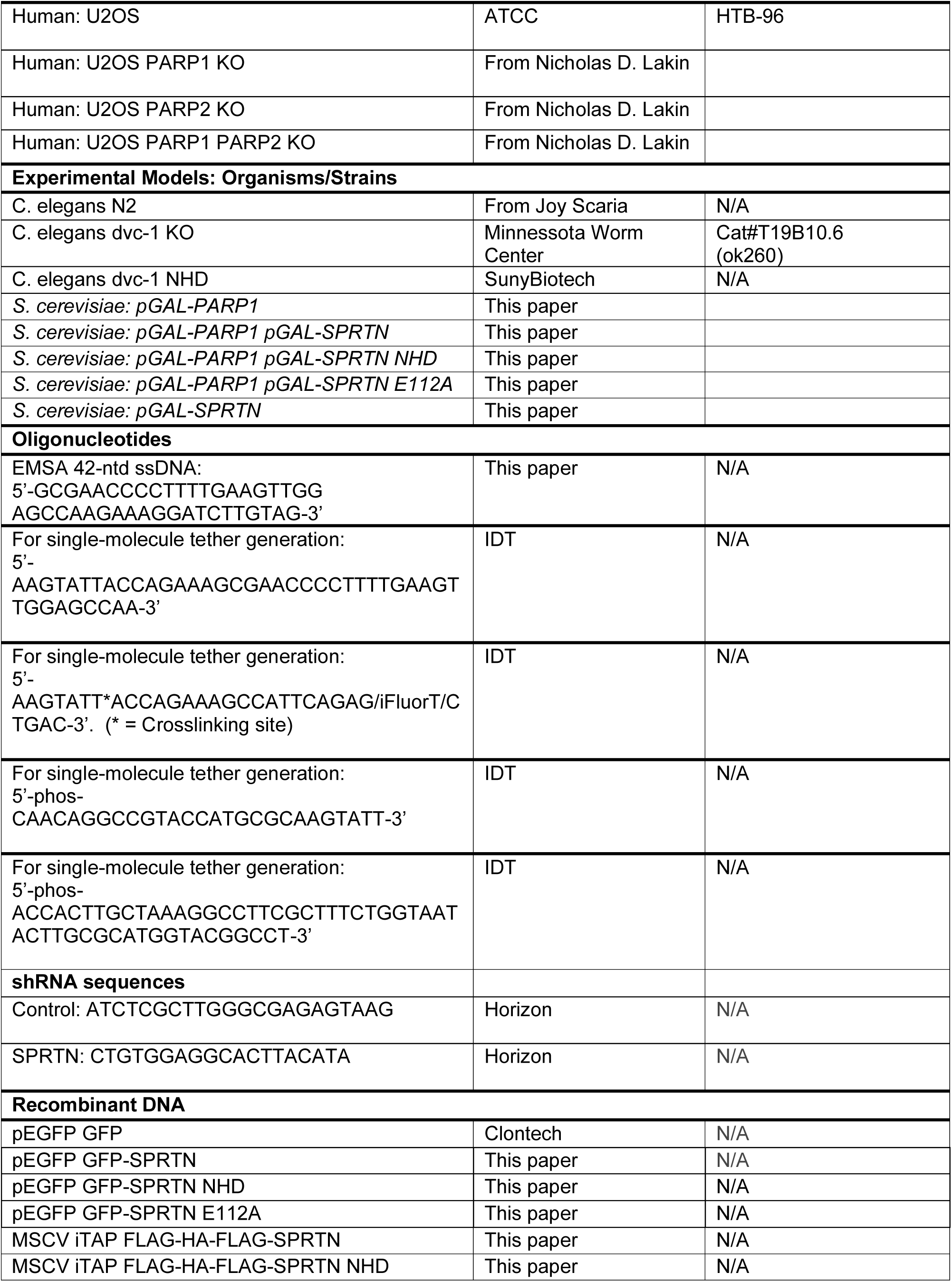
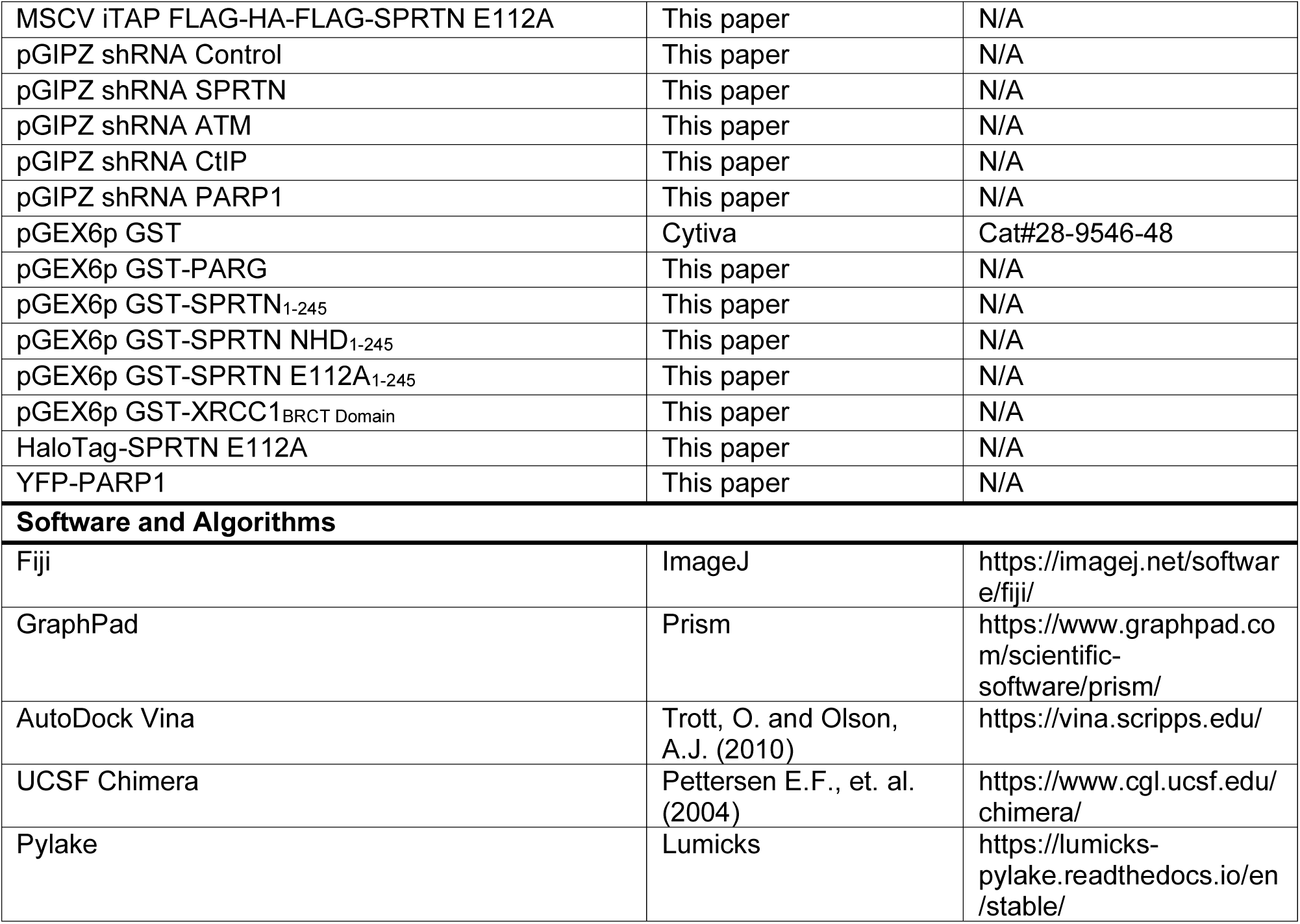

### RESOURCE AVAILABILITY

#### Lead Contact

Further information and requests for resources and reagents should be directed to the Lead Contact, Jaime Lopez-Mosqueda.

#### Materials Availability

Reagents generated in this study are available from the Lead Contact with a completed Materials Transfer Agreement.

#### Data Availability

There are no restrictions on data availability.

#### Cell lines and culture conditions

U2OS T-REx, HEK-293T, and RPE1 hTERT cell lines purchased from ATCC were cultured in DMEM with 4.5 g/L glucose, L-glutamine, and sodium pyruvate (Corning) supplemented with 10% fetal bovine serum. Cells were grown at 37°C in a humidified 5% CO_2_ incubator.

Cells were transiently transfected with Opti-MEM (Gibco) containing 1000 ng transfer gene and 5 μL TurboFect (Thermo Scientific). Media was refreshed 12 hours after transfection and experiments began 24 hours later. GFP alone and SPRTN variants cloned into pEGFP (N-terminal tag) included GFP-SPRTN WT, GFP-SPRTN NHD, GFP-SPRTN E112A, GFP-SPRTN ΔSHP, GFP-SPRTN ΔUBZ, GFP-SPRTN ΔC, and GFP-SPRTN ΔC+NLS. All plasmids were confirmed by Sanger sequencing.

For lentiviral transduction, HEK-293T cells were transfected with Opti-MEM (Gibco) containing 1000 ng psPAX2 (Addgene #12260) and 2000 ng pMD2.G (Addgene# 12259) (lentiviral packaging plasmids), 1000 ng transfer gene, and 5 μL TurboFect (Thermo Scientific). Media was refreshed 12 hours after transfection, and the lentivirus was harvested 24 hours after the addition of fresh media. Recipient cells were transduced 24 hours after plating, and the lentivirus was removed after 24 hours. Selection drug was added 48 hours post-transduction, and cells grew under selection for 10 days. All transduced cells lines were confirmed by Western blot and/or qPCR. Plasmids for stable transductions included MSCV iTAP with FLAG-HA-FLAG SPRTN WT, FLAG-HA-FLAG SPRTN NHD, and FLAG-HA-FLAG SPRTN E112A; pGIPZ with shRNA Control, shRNA PARP1, shRNA SPRTN, shRNA CtIP, and shRNA ATM.

#### Yeast Strains

W303a yeast cells were used to construct targeted gene deletions using standard yeast methods. To create yeast strains expressing PARP and FLAG-SPRTN variants under the control of the galactose inducible promoter, genes were cloned into pDONR223 (Invitrogen, Germany) using the Gateway cloning system. SPRTN variants were subsequently shuttled into pAG-306GAL-ccdB and PARP1 was shuttled into pAG-304GAL-ccdB (pAG-306GAL-ccdB and pAG-304GAL-ccdB were gifts from Susan Lindquist; Addgene plasmids #14139 and #14135). The plasmids were verified by Sanger sequencing and linearized at the URA3 or TRP1 loci and transformed into yeast strains.

### C. elegans Strains

*C. elegans* were grown and maintained at 20°C on NGM (nematode growth media) seeded with *E. coli* OP-50. Dvc-1 NHD strain was obtained from SunyBiotech, the N2 (wildtype) strain from Dr. Joy Scaria at Oklahoma State University, and the dvc-1 knock-out strain from the *Caenorhabditis* Genetics Center (University of Minnesota).

#### Bacterial Strains

*E. coli* OP-50 was used for maintenance of worms. *E. coli* Rosetta (DE3) cells were transformed with IPTG-inducible plasmids for recombinant protein purification.

#### Microirradiation

U2OS T-REx cells were plated in LabTek dishes (glass bottom) and transiently transfected with GFP-tagged constructs (pEGFP). Cells were treated with 20 μM BrdU 24 hours prior to imaging. When indicated, cells were treated with 20 μM Olaparib for 12 hours. Using the Fluoview Olympus confocal microscope, DNA damage was induced with the 405 nM UV laser (set to 100%). Images were automatically collected every 30 seconds. Signal intensity was measured using Fiji software (version 2.1.0/1.53c), and graphs were generated in GraphPad Prism (version 9.1.1).

#### Protein purification and expression

All recombinant proteins were cloned into pGEX6p1 (verified by Sanger sequencing), expressed in *E. coli* Rosetta (DE3) cells, and grown in terrific broth supplemented with 100 μg/mL ampicillin. Genes cloned into pGEX6p1 (N-terminal tag) included SPRTN WT_1-245_, SPRTN NHD_1-245_, SPRTN E112A_1-245_, SPRTN NUDIX_1-245_, GST alone, an XRCC1 fragment containing the carboxyl-terminal BRCT domain, and PARG. For SPRTN purifications, media was supplemented with 100 μM ZnCl_2_ to ensure adequate zinc for coordinating the catalytic domain. At OD600 of 0.6, protein synthesis was induced with 0.5 mM IPTG. Cultures were incubated at 16°C overnight, then pelleted, washed with 1X PBS, and stored at −80°C (1 hour or up to two weeks). Pellets were lysed in 500 mM NaCl and PBS, then incubated with lysozyme (20 minutes). Lysate was sonicated two times at 20% amplitude for 3 minutes (1 second on, 1 second off). Large pellets were sonicated four times. Cellular debris was pelleted, and supernatant was incubated with glutathione sepharose beads (GE Healthcare) (4°C rocking, 2 hours). Beads were washed four times with 500 mM NaCl and PBS. Beads were resuspended in 1X PBS. Samples were either eluted with 10 mM glutathione (Sigma, reduced form, 63H1025) in 50 mM Tris-HCl pH 8.0 or the GST tag was cleaved off with 3C PreScission protease (Sigma-Aldrich).

#### GST pull-downs

PAR was generated by incubating 0.1 mM NAD^+^, 0.1 unit/μL PARP1, and 100 ng/μL Dpn1 treated plasmid (single strand breaks; Thermo Scientific) at 37°C (30 minutes). The reaction was treated with 25 U benzonase (EMD Millipore) to degrade the DNA and 1 U ProteinaseK (Thermo Scientific) to degrade PARP1. Proteinase K was heat inactivated at 80°C. Purified recombinant GST-tagged proteins were incubated with PAR, rotating at 4°C (2 hours). Samples were washed three times with PBS, boiled, and analyzed by Ponceau stain (Acros Organics) and immunoblotting.

Purified recombinant GST-tagged proteins were also incubated with HEK-293T whole cell extract. HEK-293T cells were left untreated or treated with 20 μM PARGi (PDD 00017273: Tocris) for 12 hours. Cells were lysed in 50 mM Tris pH 8, 150 mM NaCl, 10% glycerol, 0.1% NP40, 2 mM EDTA, 1 mM PMSF, 1X protease inhibitors, and 1 mM DTT. Once sonicated and clarified, whole cell extract was incubated with purified recombinant GST-tagged proteins (rotating at 4°C, 2 hours). Samples were washed three times with PBS, boiled, and analyzed by Ponceau stain (Acros Organics) and immunoblotting.

#### Immunoblotting and Immunoprecipitation

To confirm cell lines, cells were harvested and lysed in 1X Laemmli buffer (0.031M Tris pH6.8, 1% SDS, 2.5% beta-mercaptoethanol, 5% glycerol, 0.125 mg/mL bromophenol blue). Samples were run through a polyacrylamide gel and analyzed by immunoblotting.

For immunoprecipitation, cells were harvested and lysed in 50 mM Tris-HCl pH 8.0, 150 mM NaCl, 0.1% NP40, 10% glycerol, 2 mM EDTA, 1 mM PMSF, 1 mM DTT, 1X protease inhibitors. FLAG M2 affinity beads (Sigma, SLBW2732) were used per manufacturer’s instruction to precipitate FLAG-tagged proteins. GFP-Trap-A beads (Chromotek, 71214001A) were used per manufacturer’s instruction to precipitate GFP-tagged proteins. Samples were run through a polyacrylamide gel and analyzed by immunoblotting.

To induce ADP-ribosylation in response to damage, cells were treated with 5mM H_2_O_2_ for 15 minutes at 37°C, then immediately lysed.

#### Immunofluorescence

Following microirradiation, cells were immediately fixed with methanol MES. Cells were stained with anti-GFP (Abcam, ab290, 1:500) and anti-γ-H2A.X S139 (Cell Signaling, 2577S, 1:500).

Top1cc immunofluorescence was performed as described^45^. Briefly, cells were grown on sterile coverslips and treated with 10 μM CPT (30 minutes), with 1 hour recovery or no recovery. Cells were fixed with 4% paraformaldehyde in PBS (4°C, 15 minutes) and permeabilized with 0.25% Triton X-100 in PBS (4°C, 15 minutes). Cells were treated with 1% SDS (20°C, 5 minutes), then washed five times with 0.1% BSA and 0.1% Triton X-100 in PBS. Cells were blocked in 10% nonfat powdered milk in 150 mM NaCl and 10 mM Tris-HCl pH 7.4 (20°C, 1 hour). Cells were washed once with PBS. Primary antibody (anti-Topoisomerase 1-DNA covalent complex, EMD Millipore, MABE1084, 1:100) in 5% BSA/PBS was incubated overnight (4°C). Cells were washed 6 times with 0.1% tween-20 in PBS. Secondary antibody (Donkey-anti-mouse Alexa Fluor Plus 647, Invitrogen, 1:500) in 5% BSA/PBS was incubated for 1 hour (20°C). Cells were washed 6 times with 0.1% tween-20 in PBS, with DAPI (MPbio,1 μg/mL) added to the fifth wash. Coverslips were mounted with Mowiol mounting media. Images were captured using the Fluoview Olympus confocal microscope.

#### HHpred bioinformatics and alignment

HHpred server (https://toolkit.tuebingen.mpg.de/tools/hhpred) for protein homology detection and structure prediction was used to detect functional domains in SPRTN^53^. This analysis picked up a Nudix hydrolase domain.

#### BLI measurements

BLI measurements were performed on an OctetRed system (ForteBio). Streptavidin-biosensors were loaded by placing them in biotinylated mono-ADPr, di-ADPr and tri-ADPr, 3’-biotinylated ssDNA (dA_3_), and 5’-biotinylated ssDNA (dA_3_) (150 µM) solutions and equilibrated in buffer (25 mM TRIS, 100 mM NaCl, 1mM DTT, 0.05% Tween-20, 1 mg/mL BSA, 1 mg/mL dextran, pH 7.4). Subsequently, the sensors were transferred into a solution containing SPRTN E112A (10 µM) to measure the association of the analyte. Dissociation of the ADPr_n_: SPRTN complexes was measured by placing the sensor back into buffer. Recorded spectra were analyzed using the ForteBio Data Analysis software and plotted using GraphPad Prism (version 9.1.1). Dissociation constants (K_d_) for the interaction between SPRTN and ADPr could not be reliably fitted as a lack of plateau in the association phase and an incomplete dissociation indicate additional interactions occurring, such as the dimerization of SPRTN.

### *In vitro* PAR degradation assay

PAR was synthesized as described above, but without proteinase K treatment to preserve ADP-ribosylated PARP1. In a 30 μL reaction, 1 μg of GST-tagged protein was incubated with 5 μL PAR in 50 mM Tris-HCl pH 8.0, 2 mM MgCl2, and 50 mM NaCl (30 °C, 1 or 12 hours). Reactions were stopped with 4X Laemmli buffer and boiled. Samples were run through a polyacrylamide gel and analyzed by immunoblotting.

#### SPRTN NUDIX modeling

Molecular docking was carried out to evaluate possible energy of interactions, hydrogen bonds, non-hydrogen bonds, and binding mode of di-ADP-ribose (di-ADPr) ligand with SPRTN (PDB: 6MDX). The docking studies were performed using the Autodock vina platform and visualized with UCSF chimera. In docking, SPRTN was considered semi-rigid while di-ADPr was flexible. To perform suitable docking for di-ADPr, we set the search space box parameters on 12.85–25.29–17.14 Å (direction, x, y, and z), centered at (6, 4.41, and 2.33) Å for SPRTN. The final docked conformations were ranked based on binding energy (ΔG) results, where the most favorable binding conformations had the lowest free energy of −11.4 with a RMSD of 1.336.

#### EMSA (Electrophoretic Mobility Shift Assay)

SPRTN-DNA binding was assayed as previously described^43^. Briefly, to assess DNA binding, 5 μM SPRTN was incubated in 20 mL reactions with 0.025 μM IR700 labeled 42-nt ssDNA (5’-GCGAACCCCTTTTGAAGTTGGAGCCAAGAAAGGATCTTGTAG-3’) in 25 mM Tris-HCl pH 7.5, 100 mM NaCl, and 1 mM DTT (ice for 1 hour). Increasing concentrations of PAR were added to the reactions. Samples ran through a non-SDS-PAGE 6% polyacrylamide gel with TBE buffer at room temperature. Samples were visualized with the Odyssey CLx Imaging System (Li-COR).

#### DCV DNA-protein crosslinked duplex generation

Oligonucleotides that were to be crosslinked were resuspended to 100 µM with TE Buffer. To crosslink, 1000 pmols of DCV protein were added to 100 pmols of oligo and reaction buffer (1 mM MnCl_2_, 1 mM MgCl_2_, 50 mM NaCl, 50 mM HEPES). Reaction was placed on a heat block set to 37°C for 30 minutes. Crosslinked oligonucleotide and complementary oligonucleotides were diluted to 10 µM. 50 pmols of the crosslinked oligonucleotide and 75 pmols of both the complementary top and bottom oligonucleotide were diluted in 100mM KCl, 10 mM Tris, pH 8.0 buffer and incubated at 95°C for 10 minutes. Then the heat block was turned off and gradually cooled to room temperature. Then 1 µL of the 25 nM annealed oligonucleotides were used as a substrate to LUMICKS DNA tethering kit per manufacturer’s instructions. The reaction was performed overnight and then stored at 4°C for short term, and for long term separated into single-use aliquots and frozen at −20°C. To observe ligation of the arms onto the substrate, we ran a sample on a 0.8% agarose gel at 90V for an hour and a half. Gel was scanned for the fluorescent (FAM & Atto647N) tags on the DNA and an ethidium bromide stain.

#### Cell lines

U2OS cells are cultured 5% oxygen in Dulbecco modified Eagle’s medium (DMEM) supplemented with 4.5 g/I glucose, 10% fetal bovine serum (Gibco), 5% penicillin/streptavidin (Life Technologies). In order to get the transient overexpression of YFP-tagged PARP1 and Halo-tagged catalytically dead SPRTN, 4 µg of plasmid per 4 million cells was used to transfect using the lipofectamine 2000 reagent and protocol for 24 hours. Cells expressing Halotags fusions were treated 100 nM of fluorescent Halotag ligand for 30 min at 37°C (Janelia Fluor® 552 HaloTag® Ligand from Dr. Luke Lavis Laboratory, Janelia Research Campus).

#### Nuclear extraction

After transient transfection, nuclear extraction was performed using a nuclear extraction kit from Abcam (ab113474), following the Abcam kit protocol. After the extraction, the tubes were aliquoted and flash-frozen in liquid nitrogen before being placed in −80°C. For being used on C-Trap, nuclear extracts were thawed and diluted in buffer for experiments at a ratio of 1:10.

#### Single-molecule imaging

We imaged the DPC lesion and repair factors at the single molecule level. Within the C-Trap, it contains three-color confocal fluorescence microscope and dual-trap optical tweezers. The two optical traps move between up to five flow channels separated by laminar flow. In these experiments, only four of the flow channels were used. Substrates used in the experiment were placed into flow channels one, two, three, and four, which were filled with 1.76 μm streptavidin-coated polystyrene beads (LUMICKS), ligated DPC substrate (reaction product diluted 1:300 in PBS), Buffer (20mM HEPES, 150mM NaCl, 5% glycerol, 0.5 mg/ml BSA, 1 mM DTT, 1 mM Trolox, 10 mM NAD^+^), and nuclear extracts diluted 1:10 in buffer without NAD^+^, expressing YFP-PARP1 and HaloTag-JF552-SPRTN-E112A. These were flowed through the microfluidic cell at 0.3 bar to maintain the laminar flow. In channel one of the flow cell, the polystyrene beads were first caught by both optical traps. The traps were then moved to channel two to capture the DNA. In order to capture the DNA, the bead caught in trap 2 is held at a constant position while the bead in trap 1 is kept parallel to trap 2 but is moved upstream and downstream of the flow. To tell if a single DNA tether was captured between the beads, a force-distance curve was compared to an extensible Worm-Like Chain model for a 12 kb DNA substrate. If force increases at the appropriate distance, then that confirmed that a DNA was captured. The traps were then moved to channel three to be washed by the buffer. At the same time, channel four was opened to flow in the extract into the flow cell at 0.3 bar for at least 10 seconds. After fresh nuclear extract was flowed in, the flow is stopped and the optical traps are moved to channel four. Immediately, the force curve was re-zeroed, Atto647N marker on the strand is photobleached for 20 seconds, and then the force of the DNA is moved to 10 pN for data collection.

#### Confocal imaging

To image the proteins used on the C-Trap, fluorophores on the proteins were excited with the laser that was closet to its maximum excitation wavelength. Halo-JF552-SPRTN-E112A were excited with a 561 nm laser and collected 575-625 nm band pass and YFP-PARP1 was excited with a 488 nm laser and emission collected in a 500-550 nm band pass filter. Data was collected with a 1.2 NA 60x water emersion objective and the photons were measured with single-photon avalanche photodiode detectors. The lasers of the C-Trap were set to 5% power and scanned at 100 frames per second with 0.1 msec of exposure for each pixel of size 100 nm.

#### C-Trap data analysis

Kymographs that were collected were exported and analyzed custom software by LUMICKS (Pylake). To visualize the kymographs and 2D scans, the utility C-Trap .h5 Visualization GUI (2020) by John Watters was used as it was downloaded from harbor.lumicks.com as well as Lakeview as it was downloaded from LUMICKS. Data containing both the DNA substrate and the streptavidin-coated polystyrene beads had the pixels at the end each bead defined to determine the start and end of the DNA strand. Line tracking of each event was performed using a custom script from LUMICKS that performs a Gaussian fit over the line intensity and connecting the time points to form a line using previous line tracking algorithms. For PARP1 with the YFP fluorophore, events tend to blink for periods up to two seconds which can cause a single event to seem as two separate events. To address this issue with tracking, lines that were tracked at the same position (<100 nm) with off times less than 2 secs were manually connected using a feature of the LUMICKS software. After tracking the lines, the position and time data for each line was used to determine each line’s duration, the number of lines per minute, and the average position of each line. Data from the kymographs are sorted onto a CRTD graph which is fitted on to a one-phase decay equation:

*Y = (Y0 − Plateau) * exp(−K * X) + Plateau*

Where Y0 is the Y value is at X (seconds) equals zero, plateau is the maximum Y value, and K is the rate constant which is the reciprocal of the X-axis in seconds.

#### Yeast spot assays

Yeast cells were spotted as described ^54^. Briefly, yeast cells were grown overnight in LEU2 dropout media. 0.2 OD600 of cells were collected and diluted in five-fold serial dilutions. Cells were spotted on LEU2 dropout plates containing 2 μg/mL doxycycline and/or 5 μM CPT. Yeast plates were incubated at 30 °C for 96 hr. For galactose-inducible FLAG-SPRTN expression, yeast cells were grown as above before spotting on complete plates with 2% glucose with or without 5 μM CPT or 2% galactose with or without 5 μM CPT.

### C. elegans genotyping

*C. elegans* were genotyped through PCR and restriction enzyme digest. Briefly, five adult worms were lysed in 100mM KCl, 20 mM Tris pH 8.2, 5 mM MgCl_2_, 0.9% NP40, 0.9% Tween-20, and 20 μg proteinase K. Worms were frozen at −80 °C (5 minutes) then immediately run through a thermocycler at 60 °C (1 hour) and 95 °C (15 minutes). A Taq polymerase PCR mix was made with primers to amplify dvc-1. Samples were run on standard thermocycler settings for 35 rounds of amplification. PCR fragments were digested with the restriction enzyme PstI to differentiate between N2 (wildtype), dvc-1 NHD, and dvc-1 knockout sequences.

### *C. elegans* embryo survival assay

Embryo survival was analyzed as described^52, 55^. Briefly, ten L4 staged worms of each genotype were grown on NGM plates enriched with 1 μM camptothecin or 50 μM Etoposide and OP50 (20°C, 16 hours). Worms were transferred to fresh plates (no drug) to lay eggs (20°C, 4 hours). After 4 hours, L4 worms were removed, and embryos were incubated for 24 hours (20°C) to allow for hatching. Surviving embryos were counted. Survival rates after three replicates (total n = 30 L4 worms per genotype) were determined by comparing drug treated embryo survival to no treatment survival.

### *C. elegans* longevity measurement

*C. elegans* lifespan was analyzed as described^56^. Briefly, worms were synchronized using bleaching methods (1% bleach/ 1N NaOH) to isolate eggs. Eggs were allowed three days to hatch on NGM/OP50 plates. Once at the L4 stage, 60 L4 worms were moved to NGM/OP50 plates containing 50 μM FUDR, to prevent egg hatching, and 100 μg/mL ampicillin, to prevent bacterial contamination (incubated at 20°C). The number of living worms was recorded three times a week and daily toward the end of the experiment. Counting ceased once all worms died. Graphical analysis was calculated using a Kaplan-Meier curve. For each genotype, 60 worms were analyzed across two independent experiments.

## QUANTIFICATION AND STATISTICAL ANALYSIS

Quantification and statistical analysis were performed using GraphPad Prism. Parameters, sample size, and replicates are listed in Method Details.

---

**Figure S1.** PARP inhibitors block ADP-ribosylation. (A) Immunoblot of U2OS T-REx cells treated with H_2_O_2_ (5 mM, 15 minutes) to induced genome-wide DNA damage and PARylation. Olaparib (20 μM, 12hours) and Talazoparib (1 μM, 12 hours) block H_2_O_2_-induced PARylation. Phosphorylated-Chk1 is a marker for H_2_O_2_-induced damage. (B) Microirradiation of U2OS T-REx wild-type or PARP1 knockout cells transiently expressing GFP-SPRTN and sensitized with BrdU. Insets indicate time (min:s); dashed lines mark the microirradiation path. (C) Immunoblot of RPE1 cells treated with H_2_O_2_ (5 mM, 15 minutes) to induced genome-wide DNA damage and PARylation. Confirms PARP1 and/or PARP2 knock-outs. (D) Microirradiation of U2OS T-REx cells transiently expressing GFP-SPRTN ΔC+NLS and sensitized with BrdU. Insets indicate time (min:s); dashed lines mark the microirradiation path. (E) Microirradiation of U2OS T-REx cells transiently expressing GFP-Tdp1 or GFP-Tdp1_1–164_ and sensitized with BrdU, with or without Olaparib treatment (20 μM, 12 h). Insets indicate time (min:s); dashed lines mark the microirradiation path. (F) Graph represents quantification of Tdp1 recruitment in 20 cells. Error bars represent SEM.

**Figure S2.** SPRTN preferentially binds ADP-ribose. (A) Chemical structures of ADP-ribose and a deoxyadenosine monophosphate. (B) Structural model of SPRTN (PDB: 6MDX) docked with di-ADP-ribose. Inset highlights mutations within the NHD predicted to disrupt PAR binding. (C) RNA degradation assays. In vitro reactions demonstrating RNA stability in the presence of SPRTN variants. RNase A serves as the positive control. (D) EMSA of SPRTN binding to an IF700 fluorescently labeled 42-nucleotide ssDNA fragment, while SPRTN^NUDIX^ does not bind to the ssDNA fragment. (E) In vitro SPRTN cleavage assay. SPRTN was incubated with single-stranded DNA or mono-, di- or tri-ADP-ribose (Representative of three independent experiments). (F) Immunoblot confirms doxycycline-induced expressing of FLAG-HA-FLAG SPRTN variants (shRNA resistant). (G) U2OS T-REx cells were transiently transfected with GFP constructs. GFP constructs were immunoprecipitated and analyzed by immunoblotting. p97 immunoprecipitation indicates SPRTN is a properly folded, as it can bind to p97 through its SHP domain (Representative of three independent experiments).

**Figure S3** | Characterization of SPRTN NHD. (A) Structural model of SPRTN (PDB: 6MDX) bound to di-ADP-ribose, illustrating the spatial separation between the catalytic cleft and the N-terminal helical domain (NHD). (B) Ligation of 6-kb LUMICKS DNA handles onto a crosslinked DNA duplex. Lane 1: 10-kb DNA ladder. Lane 2: control ligation of LUMICKS arms (used to assess experimental ligation efficiency). Lane 3: ligation product generated using the crosslinked annealed oligo (50 fmol loaded). Samples were run on a 0.8% agarose gel at 90 V for 1.5 h and stained with ethidium bromide. (C) Formation of DCV-dependent covalent DNA crosslinks assessed by SDS–PAGE. Crosslinking reactions contained non-fluorescent ssDNA (100 pmol) and DCV protein (200 pmol; 2:1 protein:oligo ratio) in 1× crosslinking buffer (50 mM HEPES–KOH pH 8.0, 50 mM NaCl, 1 mM MgCl₂, 1 mM MnCl₂). Lane 1: molecular weight ladder. Lane 3: DCV protein alone (200pmol). Lane 5: ssDNA crosslinked to DCV (100 pmol). Lane 7: in-house non-fluorescent ssDNA crosslinked to DCV (100 pmol). Lanes 2, 4, and 6 were left empty. Samples were resolved at 100 V for 1.5 h.

**Figure S4.** SPRTN reconstitution in SPRTN-depleted cells. (A) U2OS T-REx shRNA control and shRNA SPRTN cells analyzed by immunoblotting. (B) Immunoblotting of U2OS T-REx cells expressing an shRNA targeting SPRTN with reconstituted, doxycycline inducible FLAG-HA-FLAG-SPRTN variants. (C) EMSA demonstrating SPRTN^WT^ and SPRTN^PAR^ bind an IF700 fluorescently labeled 42-nucleotide ssDNA fragment, while SPRTN^DNA^ binds less efficiently. Coomassie stain confirms increasing SPRTN protein concentration.

**Figure S5.** SPRTN and PARP1 expression in yeast and C. elegans genotyping. (A) Wss1 degron system analyzed by immunoblotting. (B) SPRTN and PARP1 expression in yeast. Yeast strains analyzed by immunoblotting. PARP1 and SPRTN variants are regulated by the galactose promoter. Dextrose represses expression of genes under the galactose promoter. (C) PCR amplification and restriction enzyme digestion of *C. elegans* genotypes - N2 (wildtype), dvc-1^NHD^ mutant, and dvc-1 knockout. NHD mutations create a PstI restriction site that serves as a diagnostic genotyping tool. *C. elegans* dvc-1 DNA sequences are shown, with primers in yellow and the PstI restriction enzyme site in red.

## REFERENCES

1. Hoeijmakers JH. DNA damage, aging, and cancer. N Engl J Med 361, 1475–1485 (2009).

2. Elledge SJ. Cell cycle checkpoints: preventing an identity crisis. Science 274, 1664–1672 (1996).

3. Jackson SP, Bartek J. The DNA-damage response in human biology and disease. Nature 461, 1071–1078 (2009).

4. Lopez-Otin C, Blasco MA, Partridge L, Serrano M, Kroemer G. The hallmarks of aging. Cell 153, 1194–1217 (2013).

5. Duxin JP, Dewar JM, Yardimci H, Walter JC. Repair of a DNA-protein crosslink by replication-coupled proteolysis. Cell 159, 346–357 (2014).

6. Ruijs MW, van Andel RN, Oshima J, Madan K, Nieuwint AW, Aalfs CM. Atypical progeroid syndrome: an unknown helicase gene defect? Am J Med Genet A **116A**, 295–299 (2003).

7. Lessel D, et al. Mutations in SPRTN cause early onset hepatocellular carcinoma, genomic instability and progeroid features. Nat Genet 46, 1239–1244 (2014).

8. Vaz B, et al. Metalloprotease SPRTN/DVC1 Orchestrates Replication-Coupled DNA-Protein Crosslink Repair. Mol Cell 64, 704–719 (2016).

9. Stingele J, et al. Mechanism and Regulation of DNA-Protein Crosslink Repair by the DNA-Dependent Metalloprotease SPRTN. Mol Cell 64, 688–703 (2016).

10. Lopez-Mosqueda J, et al. SPRTN is a mammalian DNA-binding metalloprotease that resolves DNA-protein crosslinks. Elife 5, (2016).

11. Painter RB, Young BR. Radiosensitivity in ataxia-telangiectasia: a new explanation. Proc Natl Acad Sci U S A 77, 7315–7317 (1980).

12. Ciccia A, Elledge SJ. The DNA damage response: making it safe to play with knives. Mol Cell 40, 179–204 (2010).

13. Bergink S, Jentsch S. Principles of ubiquitin and SUMO modifications in DNA repair. Nature 458, 461–467 (2009).

14. Harper JW, Elledge SJ. The DNA damage response: ten years after. Mol Cell 28, 739–745 (2007).

15. Kleine H, Luscher B. Learning how to read ADP-ribosylation. Cell 139, 17–19 (2009).

16. Ruggiano A, et al. The protease SPRTN and SUMOylation coordinate DNA-protein crosslink repair to prevent genome instability. Cell Rep 37, 110080 (2021).

17. Serbyn N, et al. SUMO orchestrates multiple alternative DNA-protein crosslink repair pathways. Cell Rep 37, 110034 (2021).

18. Liu JCY, et al. Mechanism and function of DNA replication-independent DNA-protein crosslink repair via the SUMO-RNF4 pathway. EMBO J, e107413 (2021).

19. Borgermann N, et al. SUMOylation promotes protective responses to DNA-protein crosslinks. EMBO J 38, (2019).

20. Schellenberg MJ, et al. Ubiquitin stimulated reversal of topoisomerase 2 DNA-protein crosslinks by TDP2. Nucleic Acids Res 48, 6310–6325 (2020).

21. Larsen NB, et al. Replication-Coupled DNA-Protein Crosslink Repair by SPRTN and the Proteasome in Xenopus Egg Extracts. Mol Cell, (2018).

22. Sun Y, Miller Jenkins LM, Su YP, Nitiss KC, Nitiss JL, Pommier Y. A conserved SUMO pathway repairs topoisomerase DNA-protein cross-links by engaging ubiquitin-mediated proteasomal degradation. Sci Adv 6, (2020).

23. Nakano T, Xu X, Salem AMH, Shoulkamy MI, Ide H. Radiation-induced DNA-protein cross-links: Mechanisms and biological significance. Free Radic Biol Med 107, 136–145 (2017).

24. Fabian Z, et al. PARP1-dependent DNA-protein crosslink repair. Nat Commun 15, 6641 (2024).

25. Shao Z, et al. Clinical PARP inhibitors do not abrogate PARP1 exchange at DNA damage sites in vivo. Nucleic Acids Res 48, 9694–9709 (2020).

26. Ronson GE, et al. PARP1 and PARP2 stabilise replication forks at base excision repair intermediates through Fbh1-dependent Rad51 regulation. Nat Commun 9, 746 (2018).

27. Centore RC, Yazinski SA, Tse A, Zou L. Spartan/C1orf124, a reader of PCNA ubiquitylation and a regulator of UV-induced DNA damage response. Mol Cell 46, 625–635 (2012).

28. Machida Y, Kim MS, Machida YJ. Spartan/C1orf124 is important to prevent UV-induced mutagenesis. Cell Cycle 11, 3395–3402 (2012).

29. Davis EJ, Lachaud C, Appleton P, Macartney TJ, Nathke I, Rouse J. DVC1 (C1orf124) recruits the p97 protein segregase to sites of DNA damage. Nat Struct Mol Biol 19, 1093–1100 (2012).

30. Mosbech A, et al. DVC1 (C1orf124) is a DNA damage-targeting p97 adaptor that promotes ubiquitin-dependent responses to replication blocks. Nat Struct Mol Biol 19, 1084–1092 (2012).

31. Pouliot JJ, Yao KC, Robertson CA, Nash HA. Yeast gene for a Tyr-DNA phosphodiesterase that repairs topoisomerase I complexes. Science 286, 552–555 (1999).

32. Das BB, et al. PARP1-TDP1 coupling for the repair of topoisomerase I-induced DNA damage. Nucleic Acids Res 42, 4435–4449 (2014).

33. Kliza KW, Liu Q, Roosenboom LWM, Jansen P, Filippov DV, Vermeulen M. Reading ADP-ribosylation signaling using chemical biology and interaction proteomics. Mol Cell 81, 4552–4567 e4558 (2021).

34. Ahel I, et al. Poly(ADP-ribose)-binding zinc finger motifs in DNA repair/checkpoint proteins. Nature 451, 81–85 (2008).

35. Wang Z, et al. Recognition of the iso-ADP-ribose moiety in poly(ADP-ribose) by WWE domains suggests a general mechanism for poly(ADP-ribosyl)ation-dependent ubiquitination. Genes Dev 26, 235–240 (2012).

36. Zimmermann L, et al. A Completely Reimplemented MPI Bioinformatics Toolkit with a New HHpred Server at its Core. J Mol Biol 430, 2237–2243 (2018).

37. Gabler F, et al. Protein Sequence Analysis Using the MPI Bioinformatics Toolkit. Curr Protoc Bioinformatics 72, e108 (2020).

38. Li J, et al. A conserved NAD(+) binding pocket that regulates protein-protein interactions during aging. Science 355, 1312–1317 (2017).

39. Daniels CM, Thirawatananond P, Ong SE, Gabelli SB, Leung AK. Nudix hydrolases degrade protein-conjugated ADP-ribose. Sci Rep 5, 18271 (2015).

40. Koonin EV. A common set of conserved motifs in a vast variety of putative nucleic acid-dependent ATPases including MCM proteins involved in the initiation of eukaryotic DNA replication. Nucleic Acids Res 21, 2541–2547 (1993).

41. Bessman MJ, Frick DN, O’Handley SF. The MutT proteins or “Nudix” hydrolases, a family of versatile, widely distributed, “housecleaning” enzymes. J Biol Chem 271, 25059–25062 (1996).

42. Palazzo L, et al. Processing of protein ADP-ribosylation by Nudix hydrolases. Biochem J 468, 293–301 (2015).

43. Li F, Raczynska JE, Chen Z, Yu H. Structural Insight into DNA-Dependent Activation of Human Metalloprotease Spartan. Cell Rep 26, 3336–3346 e3334 (2019).

44. Chandler M, de la Cruz F, Dyda F, Hickman AB, Moncalian G, Ton-Hoang B. Breaking and joining single-stranded DNA: the HUH endonuclease superfamily. Nat Rev Microbiol 11, 525–538 (2013).

45. Patel AG, et al. Immunodetection of human topoisomerase I-DNA covalent complexes. Nucleic Acids Res 44, 2816–2826 (2016).

46. Redinbo MR, Stewart L, Kuhn P, Champoux JJ, Hol WG. Crystal structures of human topoisomerase I in covalent and noncovalent complexes with DNA. Science 279, 1504–1513 (1998).

47. Stingele J, Schwarz MS, Bloemeke N, Wolf PG, Jentsch S. A DNA-dependent protease involved in DNA-protein crosslink repair. Cell 158, 327–338 (2014).

48. Balakirev MY, et al. Wss1 metalloprotease partners with Cdc48/Doa1 in processing genotoxic SUMO conjugates. Elife 4, (2015).

49. Pommier Y, Huang SY, Gao R, Das BB, Murai J, Marchand C. Tyrosyl-DNA-phosphodiesterases (TDP1 and TDP2). DNA Repair (Amst) 19, 114–129 (2014).

50. La Ferla M, et al. Expression of human poly (ADP-ribose) polymerase 1 in Saccharomyces cerevisiae: Effect on survival, homologous recombination and identification of genes involved in intracellular localization. Mutat Res 774, 14–24 (2015).

51. Gnanasundram SV, Kos M. Fast protein-depletion system utilizing tetracycline repressible promoter and N-end rule in yeast. Mol Biol Cell 26, 762–768 (2015).

52. Dokshin GA, et al. GCNA Interacts with Spartan and Topoisomerase II to Regulate Genome Stability. Dev Cell 52, 53–68 e56 (2020).

53. Soding J, Biegert A, Lupas AN. The HHpred interactive server for protein homology detection and structure prediction. Nucleic Acids Res 33, W244–248 (2005).

54. Lopez-Mosqueda J, Maas NL, Jonsson ZO, Defazio-Eli LG, Wohlschlegel J, Toczyski DP. Damage-induced phosphorylation of Sld3 is important to block late origin firing. Nature 467, 479–483 (2010).

55. MacQueen AJ, Villeneuve AM. Nuclear reorganization and homologous chromosome pairing during meiotic prophase require C. elegans chk-2. Genes Dev 15, 1674–1687 (2001).

56. Sutphin GL, Kaeberlein M. Measuring Caenorhabditis elegans life span on solid media. J Vis Exp, (2009).

