## Supplementary figures and images for "PARP1 recruits SPRTN to DNA-protein crosslinks through a conserved poly-ADP-ribose binding domain"

### Supplemental Figure 1

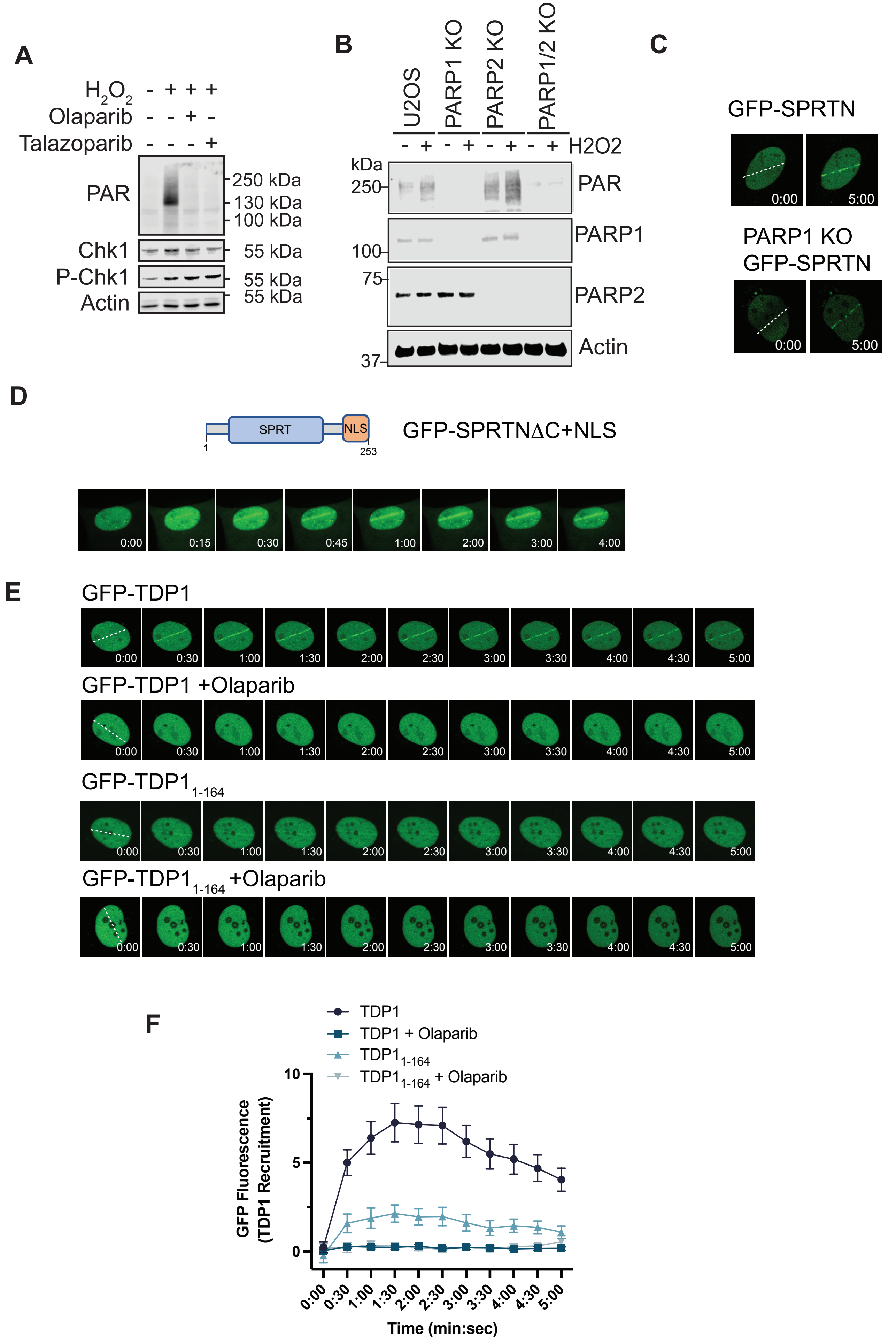

### Supplemental Figure 2

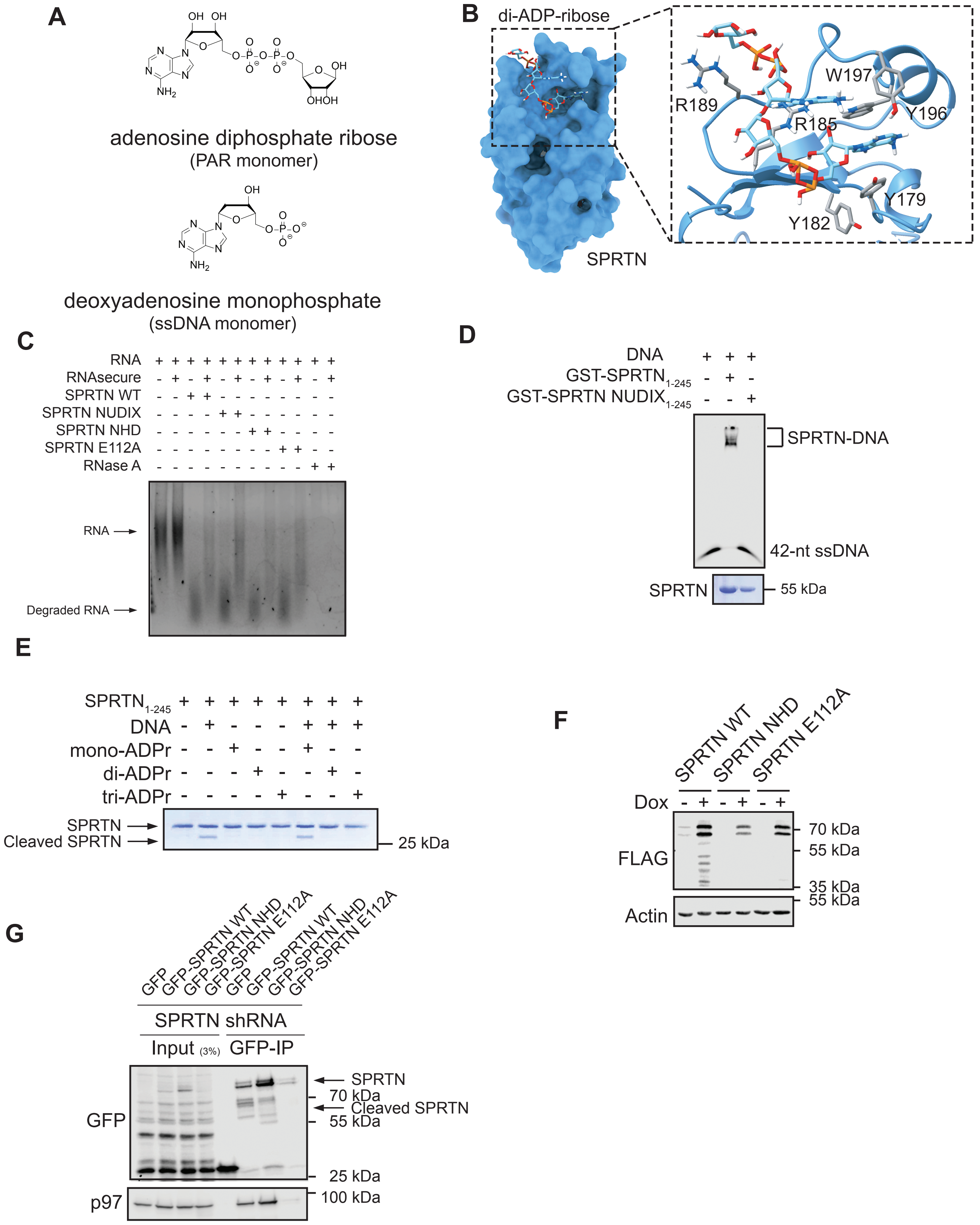

### Supplemental Figure 3

#### Figure Sup 3

A

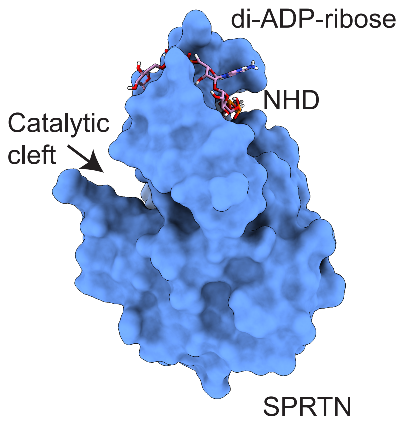


B

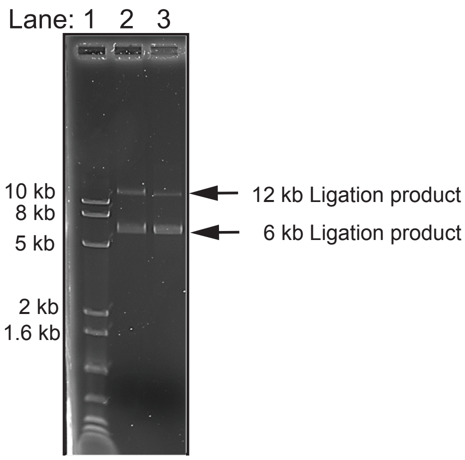


C

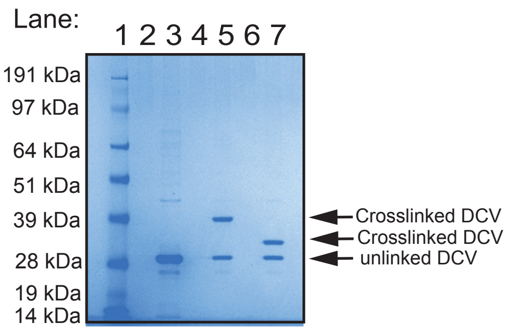

### Supplemental Figure 4

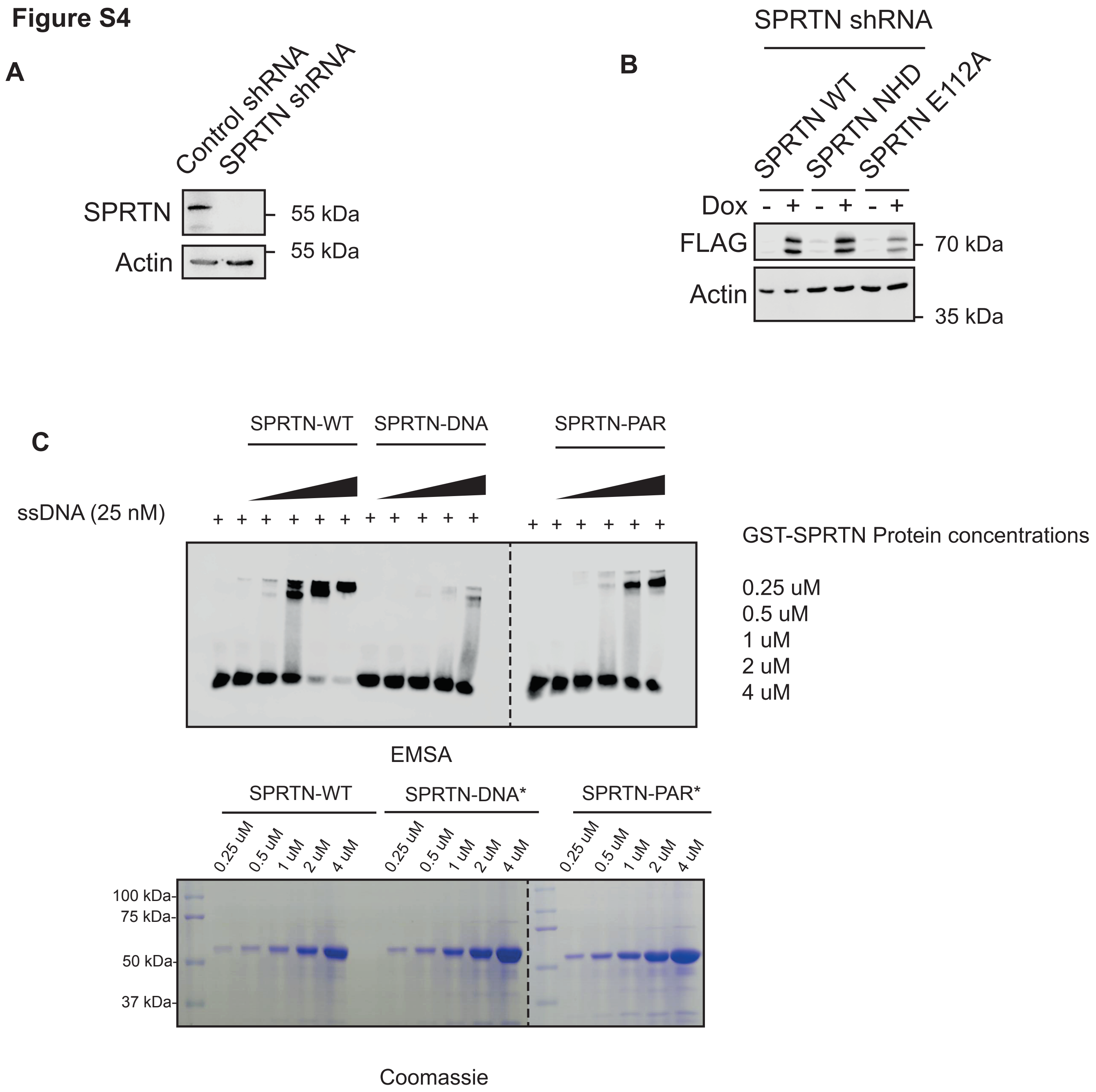
