## Supplemental Figure 5 for "PARP1 recruits SPRTN to DNA-protein crosslinks through a conserved poly-ADP-ribose binding domain"

**Figure S5**

**A**

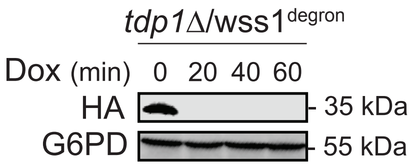


**B**

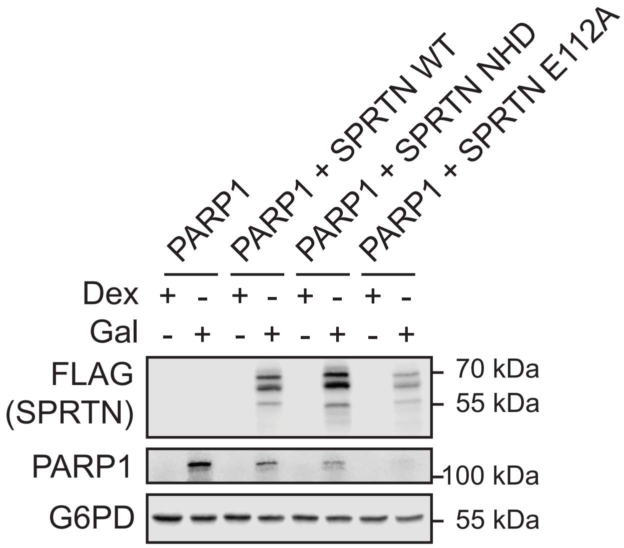


**C**

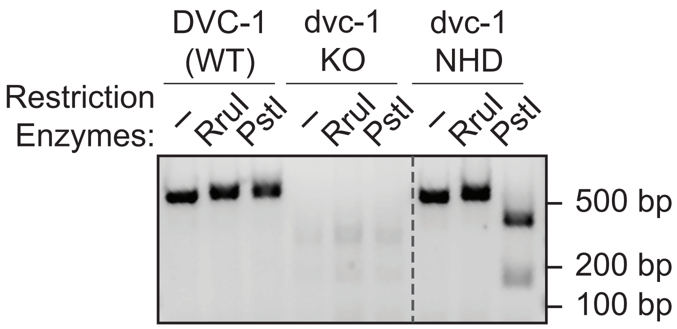


DVC-1 (WT): Dvc-1 coding sequence

ATGTGCCGGAATATGTTCATATGAAATTCGTGGTGGTCGTGGAGGACTTTGTTCGATTCGGC
TCAGTAAACCCCTGTTGACATTAAGACCGAGAAGTGATCTTGTTGAGACCCTGCTGCATGAA
ATGATCCACGCATATCTTTTTGTTAAAGAACGAAATAGAGATCGTGATGGTCACGGCCCACAA
TTCCAGGCCCATATGCACAGAATTAACCAAGCTGGTGGAACTAATATCACGATTTATCACAGT
TTTCATGACGAAGTTCGGCTTTACAAACAACATTGGTGGAGATGTAGTGGACCGTGTAGAGA
CCGTCGCCCATTTTTTGGATATGTGAAAAGATCCTGCAATCGAGCACCCGGACCGAATGATA
GATGGTGGAGTCAACATCAGCAAAGCTGTGGAGGTTAGACTTTATTTTTCTAATAAATATTTTT
TCATTTTTTACATTTCAGGTAACTTTCTGAAAGTTAAGG

Dvc-1 NHD: dcv-1 coding sequence

ATGTGCCGGAATATGTTCATATGAAATTCGTGGTGGTCGTGGAGGACTTTGTTCGATTCGGC
TCAGTAAACCCCTGTTGACATTAAGACCGAGAAGTGATCTTGTTGAGACCCTGCTGCATGAA
ATGATCCACGCATATCTTTTTGTTAAAGAACGAAATAGAGATCGTGATGGTCACGGCCCACAA
TTCCAGGCCCATATGCACAGAATTAACCAAGCTGGTGGAACTAATATCACGATTTATCACAGT
TTTCATGACGAAGTTCGGCTTTACAAACAGCACTGGTGGAGATGTAGTGCGCCGTGTAGGG
ACCGTCGCCCAGCTGCA|GGATATGTGAAAGCATCCTGCAATCGAGCACCCGGACCGAACG
ATAGATGGTGGAGTCAACATCAGCAAAGCTGTGGAGGTTAGACTTTATTTTTCTAATAAATATT
TTTTCATTTTTTACATTTCAGGTAACTTTCTGAAAGTTAAGG

Yellow = primers
| = RE site
